# Intrinsically disordered regions navigate DNA replication licensing factors to specific genomic loci

**DOI:** 10.64898/2026.09.12.751089

**Authors:** Innesa Leonovich, Matthew W. Parker

**Affiliations:** Department of Biophysics, University of Texas Southwestern Medical Center, Dallas, TX 75235, USA

**Keywords:** ORC, Orc1, DNA replication, intrinsically disordered region, phase separation, DNA-binding proteins, genome binding

## Abstract

Sites of DNA replication initiation are determined by binding of the Origin Recognition Complex (ORC, composed of Orc1-6) to specific genomic loci. Here, we determine the mechanism underlying site-specific binding of human Orc1. We find that Orc1’s Bromo-Adjacent Homology (BAH) domain functions as an avidity element, whereas an intrinsically disordered region (IDR) guides genomic specificity. Deletion of the IDR abolishes chromatin binding and Orc1 chimeras with swapped IDRs are redirected to new genomic locations. Other replication factors, such as Cdt1 and Cdc6, also possess IDRs. However, despite their similar biochemical properties, we find that these IDRs are functionally non-equivalent. Like Orc1, the Cdt1 IDR guides loci-specific binding but targets distinct sites. Strikingly, the Cdt1 IDR, when swapped into Orc1, redirects Orc1 to Cdt1 binding sites. This establishes a key role for the Orc1 IDR in genome navigation and points to impressive levels of functional sophistication in licensing factor disordered regions.

## INTRODUCTION

Chromatin-binding proteins are abundant in the proteome and play essential roles in chromosome organization, transcriptional regulation, DNA repair, and DNA replication. Interactions with chromatin are established through DNA and histone contacts that vary in both specificity and the extent of the chromatin interface (*1*). For example, histone reader proteins target large genomic territories by binding a specific epigenetic mark (kilobase- to megabase-pair resolution), such as seen for Heterochromatin protein 1 (Hp1) binding to H3K9me3 (*2*), a signature of transcriptionally silent chromatin (*3*). Conversely, DNA-binding domains can target specific chromosomal loci (base pair resolution) by recognizing DNA consensus motifs (*4*). Such site-specific interactions are important for transcriptional regulation and the initiation of DNA replication (*5*, *6*).

Site-specific transcription factor binding is strongly influenced by the presence of sequence-specific DNA-binding domains (*7*), a structurally diverse family of proteins with varying degrees of specificity (*8*, *9*). However, it has long been recognized that DNA-binding domains only partially explain target site selection (*10–13*) and it has become increasingly apparent that, beyond cis regulatory elements, multiple additional factors influence transcription factor binding, including local DNA structure, partner protein interactions, and epigenetic information (*13*). How transcription factors interpret these complex molecular cues is an important question, and recent work highlights the role of protein intrinsically disordered regions (IDRs) as additional specificity determinants (*12*, *14–16*). Indeed, transcription factor IDRs have been shown to target specific promoters independently of their DNA-binding domains (DBDs). The mechanism(s) underlying IDR-mediated specificity is currently under investigation and may involve direct DNA contacts or multivalent, context-dependent interactions with the chromatin environment (*16*, *17*).

The selection of DNA replication initiation sites is, like transcription, mechanistically complex and, despite the broad conservation of factors, evolutionarily divergent (*18*). In eukaryotes, the Origin Recognition Complex (ORC, composed of Orc1-6) binds specific genomic sites, known as “origins”, and marks them for DNA replication initiation. ORC functions by nucleating the assembly of a Pre-Replication Complex (Pre-RC), which loads the replicative helicase, Mcm2-7, onto origin DNA (*19*, *20*). Later, in S phase, Mcm2-7 is activated to unwind duplex DNA and form a molecular platform for replisome assembly (*21*). Similar to transcription factors, budding and fission yeast ORC possess DNA-binding elements that recognize specific cis regulatory sequences (*22–24*). Interestingly, these DNA-binding elements are species-specific, as are the sequences to which they bind. In contrast to yeast, metazoan ORC lacks a sequence-specific DNA-binding domain and binds DNA promiscuously (*25*); accordingly, ORC’s *in vivo* binding sites, while specific, lack a shared DNA sequence motif (*26*, *27*). The mechanism underlying genomic site selection of metazoan ORC remains unknown.

Here, we demonstrate the mechanism of site-specific chromosome binding of human ORC. We show that two domains of Orc1, the Bromo-Adjacent Homology (BAH) domain and intrinsically disordered region (IDR), synergize to drive dynamic chromatin association. We find that the BAH domain functions as a non-specific avidity element, while the IDR drives site selection. In the absence of the IDR, Orc1 is displaced from chromatin, and swapping the Orc1 IDR for other chromatin-binding disordered regions redirects the chimeric proteins to new genomic locations. Many other Pre-RC components also have long IDRs, and we find that they are functionally non-equivalent. Specifically, although both Cdt1 and Cdc6 have IDRs with Orc1-like amino acid composition, only the Cdt1 IDR binds chromatin, and its genome binding specificity is distinct from Orc1. Notably, a chimeric Orc1 containing the Cdt1 IDR more closely resembles Cdt1’s genome binding profile than it does Orc1’s. Altogether, these studies reveal the source of metazoan ORC’s genome binding specificity and, more broadly, demonstrate that IDRs can facilitate genome navigation in the absence of traditional sequence-specific DNA binding domains.

## RESULTS

### Orc1’s folded domains are dispensable for site-specific chromosome binding

The human Origin Recognition Complex (ORC) is labile, and the dissociable complexes (Orc1, Orc2-5, and Orc6) can independently engage chromatin (*28–30*). The Orc1 subunit uniquely possesses multiple DNA and chromatin binding elements, including an N-terminal Bromo Adjacent Homology (BAH) domain connected via a long intrinsically disordered region (IDR) to an ATPases Associated with diverse cellular Activities (AAA+) and winged-helix (wH) domain (**Fig. 1A**). The Orc1 BAH domain binds dimethylated Histone H4 at lysine 20 (H4K20me2) (*31*) and the AAA+ and wH domains, when integrated into the ORC core complex, encircle DNA in an ATP-dependent fashion (*32*). The fly Orc1 IDR has additionally been shown to bind DNA in a sequence-nonspecific fashion (*33*). How Orc1’s multiple DNA and chromatin binding elements coordinately engage chromatin to facilitate genome site selection remains unknown.

**Figure 1:**
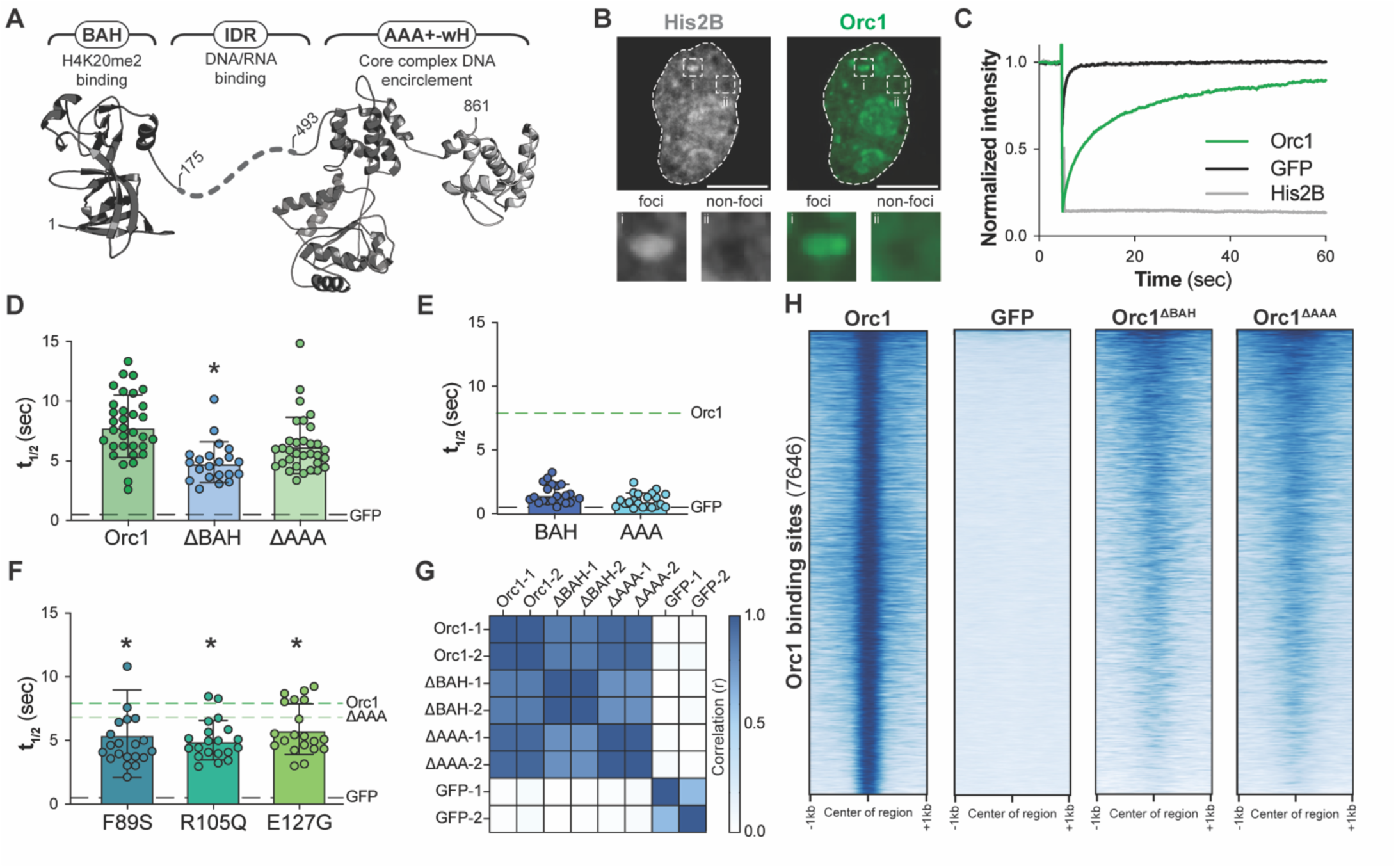
Orc1’s folded domains are dispensable for site-specific chromatin binding. (A) The architecture of Orc1 showing its multiple DNA and chromatin-binding elements. (B) Nuclear distribution of mCherry-tagged Histone H2B and GFP-tagged Orc1 in HeLa cells; scale bar = 10 µm. (C) Analysis of GFP (black line), mCherry-H2B (grey line), and GFP-Orc1 (green line) nuclear FRAP dynamics in HeLa cells. Each line represents the average of >9 individual FRAP curves. (D) To quantify chromatin binding propensity, the half-time of recovery (t_1/2_) was calculated from FRAP analyses of GFP-tagged Orc1, Orc1^ΔBAH^, and Orc1^ΔAAA^. Each marker represents a separate cell, and the mean and standard deviation are plotted. Significance was established through a one-way ANOVA test compared to full-length Orc1 (* = p < 0.05). (E) Half-time of recovery for the isolated Orc1 BAH and AAA+-wH domains. (F) Half-time of recovery for Orc1^ΔAAA^ constructs containing the indicated Meier-Gorlin Syndrome mutations in the BAH domain. (G) Correlation analysis of CUT&RUN genome coverage files for replicates of Orc1, Orc1^ΔBAH^, and Orc1^ΔAAA^. (H) Heat map representation of sequencing read intensity of GFP (second panel), Orc1^ΔBAH^ (third panel), and Orc1^ΔAAA^ (fourth panel) across Orc1 peaks (+/− 1 kb) (first panel).

To understand the mechanism of Orc1 chromatin binding, we expressed Orc1 and Orc1 sub-domain constructs in HeLa cells with a C-terminal GFP tag and used fluorescence recovery after photobleaching (FRAP) to quantify nuclear dynamics as a measure of chromatin association, as has been done previously (*34–36*). Orc1 protein levels are tightly controlled (*29*), and the full-length Orc1 transgene was expressed at near-endogenous levels (**Fig. S1A**) and exhibited the expected non-uniform, punctate distribution in the nucleus (**Fig. 1B**) (*37*). Prior work identifies these Orc1-enriched nuclear foci as heterochromatin (*35*). The FRAP dynamics of the diffusive, non-foci population of Orc1 (**Fig. 1C**, green line) indicated that Orc1 is dynamically bound to chromatin, with recovery dynamics significantly faster than those of Histone H2B (**Fig. 1C**, grey line) but substantially slower than GFP alone (**Fig. 1C**, black line). Cell fractionation and western blotting confirmed a dynamic association with chromatin, with Orc1 present in both the nuclear soluble and nuclear-bound fractions (**Fig. S1B**).

We next determined the contribution of Orc1’s folded domains to chromatin binding. Specifically, we assayed FRAP dynamics for Orc1 constructs lacking either the BAH (Orc1^ΔBAH^) or AAA+-wH domains (Orc1^ΔAAA^) and calculated the half-time of recovery (t_1/2_) to quantitatively compare their chromatin binding propensity (**Fig. 1D**). We found that removal of the BAH domain (**Fig. 1D**, ΔBAH t_1/2_ = 4.9 sec ± 1.7) significantly reduced the half-time of recovery compared to Orc1 (Orc1 t_1/2_ = 7.9 sec ± 2.6), although not to the level of the negative control (GFP t_1/2_ = 0.5 sec ± 0.6), indicating that some level of chromatin binding is retained. On the other hand, removal of the AAA+-wH domain did not significantly affect chromatin binding dynamics (**Fig. 1D**, ΔAAA t_1/2_ = 6.3 sec ± 2.4), and neither the BAH nor AAA+-wH domains on their own showed evidence of chromatin binding (**Fig. 1E**, AAA t_1/2_ = 1.0 sec ± 0.5 and BAH t_1/2_ = 1.5 sec ± 0.7). We also assayed the impact of BAH domain mutations implicated in Meier-Gorlin Syndrome (*38*), a form of primordial dwarfism. These mutations mimicked Orc1^ΔBAH^ and led to faster FRAP recovery, indicating a reduced level of chromatin binding (**Fig. 1F**, F89S t_1/2_ = 5.5 sec ± 3.4, R105Q t_1/2_ = 5.0 sec ± 1.5, E127G t_1/2_ = 5.8 sec ± 1.9).

The non-equivalent roles of the Orc1 BAH and AAA+-wH domains motivated us to investigate their contributions to site-specific chromatin binding. We therefore performed cleavage under targets and release using a nuclease (CUT&RUN) (*39*) to compare the genome-binding profiles of Orc1, Orc1^ΔBAH^, and Orc1^ΔAAA^. Consistent with prior genomics work targeting the endogenous protein, Orc1 peaks identified by CUT&RUN analysis of Flag-GFP-tagged Orc1 were enriched in H3K4me3 (**Fig. S1C**, left panel) (*27*) and were largely, though not exclusively, present in open regions of the genome (**Fig. S1C**, right panel). We observed high reproducibility between the genome coverage files of two Orc1 replicates (**Fig. 1G**, Orc1-1 and Orc1-2). Using the same CUT&RUN approach, we assessed how removal of either Orc1’s BAH or AAA+-wH domain impacts genomic localization (**Fig. 1H**). Deletion of either domain resulted in an overall reduction in signal intensity across Orc1 peaks (**Fig. 1H**). However, both Orc1^ΔBAH^ and Orc1^ΔAAA^ retained the ability to specifically localize to the same genomic regions occupied by full-length Orc1. Consistently, Pearson’s correlation analysis of Orc1, Orc1^ΔBAH^, and Orc1^ΔAAA^ genome coverage files revealed significant correlation between full-length Orc1 and the two domain deletion constructs (**Fig. 1G**). The correlation between Orc1 and Orc1^ΔBAH^ (r = 0.88) was slightly weaker than what was observed for Orc1^ΔAAA^ (r = 0.95), consistent with the BAH domains contribution to bulk chromatin binding (**Fig. 1D**). These data demonstrate that Orc1’s folded domains, and especially the BAH domain, help stabilize Orc1 at specific loci but are not strictly required for Orc1 to navigate to these sites.

### The Orc1 IDR is necessary for chromatin recruitment and genome binding specificity

The ability of Orc1 to maintain site-specific chromosome binding in the absence of its folded domains motivated us to assess the contribution of the Orc1 IDR (**Fig. 2A**, green highlighted region) to genome localization. We have previously shown that the fly Orc1 IDR binds DNA non-specifically (*33*), and we therefore assessed the DNA binding affinity of the purified human Orc1 IDR (Orc1^IDR^, residues 175-493, **Fig. 2B-C**). We found that Orc1^IDR^ bound a FITC-labeled 60-basepair oligonucleotide of random sequence (50% GC content) with high affinity (K_d_ = 81.6 ± 3 nM). Consistently, FRAP analysis in HeLa cells showed a significantly faster Orc1 recovery rate in the absence of the IDR (**Fig. 2D**, Orc1^ΔIDR^ t_1/2_= 2.5 sec ± 1.3), indicating a marked loss in chromatin binding. Removal of the IDR had a larger effect on chromatin binding than removal of either of Orc1’s folded domains (**Fig. 1D**). Although necessary, the IDR on its own was not sufficient for normal chromatin binding (**Fig. 2D**, Orc1^IDR^ t_1/2_= 3.6 sec ± 1.5). Notably, the human Orc1 IDR has an abundance of Cyclin-Dependent Kinase (CDK) phosphorylation sites (**Fig. 2E**, consensus = ‘[S/T]P’), and we asked whether regulatory phosphorylation might explain the discrepancy between the isolated IDR’s high-affinity DNA binding *in vitro* and what seems to be relatively weak chromatin binding in cells. We found that a variant that cannot be phosphorylated (Orc1^IDR-P-dead^ t_1/2_ = 4.3 sec ± 1.6, **Fig. 2F**) had a small but significant increase in recovery time compared to a phospho-mimetic mutant (Orc1^IDR-PM^ t_1/2_ = 3.1 sec ± 1.6, **Fig. 2F**). This suggests that phosphorylation negatively regulates the Orc1 IDR’s chromatin binding activity, similar to what we have observed with fly Orc1 (*40*). Interestingly, the impact of phosphorylation was significantly larger in HEK293 cells, and Orc1^IDR^ behaved more similarly to the phospho-defective variant, indicating that the basal level of Orc1 phosphorylation and the strength of the phospho-regulatory axis are cell type-dependent (**Fig. S2A**).

**Figure 2:**
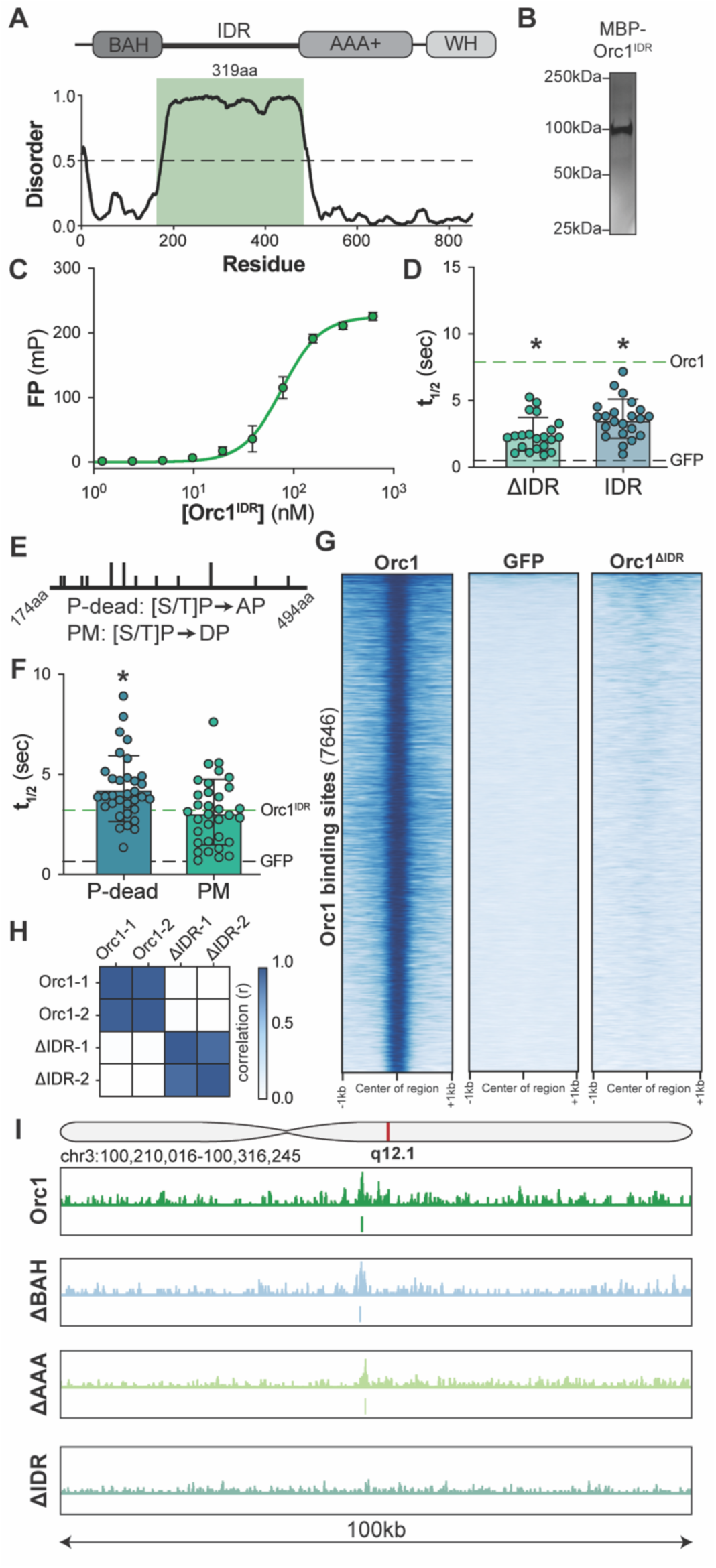
The Orc1 IDR is necessary for chromatin recruitment and genome-binding specificity. (A) Metapredict (*78*) analysis of human Orc1 highlighting an extensive region of intrinsic disorder (green highlighted region). (B) Coomassie-stained SDS-PAGE gel of purified MBP-tagged Orc1^IDR^. (C) Analysis of Orc1^IDR^’s DNA-binding affinity by fluorescence polarization (Orc1^IDR^ K_d_ = 82 ± 3 nM). (D) Half-time of recovery calculated from FRAP analyses of Orc1^ΔIDR^ and Orc1^IDR^ in HeLa cells. Each marker represents a separate cell; the mean and standard deviation are plotted. Significance was established through a one-way ANOVA test compared to full-length Orc1 (* = p < 0.05). (E) Schematic representation of phospho-site (‘[S/T]P’) distribution across the human Orc1 IDR. The indicated mutations were made to generate phospho-dead (Orc1^IDR-P-dead^) and phospho-mimetic (Orc1^IDR-PM^) variants. (F) Half-time of recovery calculated from FRAP analyses of Orc1^IDR-PM^ and Orc1^IDR-P-dead^ in HeLa cells. (G) Heat map representation of sequencing read intensity of GFP (second panel) and Orc1^ΔIDR^ (third panel) across Orc1 peaks (first panel, +/− 1 kb). (H) Correlation analysis of CUT&RUN genome coverage files for replicates of Orc1 and Orc1^ΔIDR^. (I) Genome browser view of the sequencing read intensity of full-length Orc1 and deletion constructs across a 100 kb region of chromosome 3.

We next addressed the contribution of the Orc1 IDR to genomic site selection using the same CUT&RUN approach described previously. Consistent with the observed defect in chromatin binding (**Fig. 2D**), Orc1^ΔIDR^ showed a near-complete loss of signal at sites occupied by wild-type Orc1 (**Fig. 2G**, right panel). Additionally, and in contrast to the deletion of Orc1’s folded domains (**Fig. 1G**), there was no significant correlation between Orc1 and Orc1^ΔIDR^ genome coverage files (**Fig. 2H**, r = 0.38). We further compared sequencing reads at specific chromosomal loci and observed a broad loss in Orc1^ΔIDR^ signal both across the chromosome and specifically at Orc1 peaks (**Fig. 2I**, compare top and bottom tracts). Conversely, both Orc1^ΔBAH^ and Orc1^ΔAAA^ were retained at Orc1 peaks (**Fig. 2I**, compare top three tracts). These data demonstrate that the DNA-binding Orc1 disordered region is essential for chromatin binding and, consequently, recruitment of Orc1 to specific sites.

### Swapping disordered regions retargets Orc1 to new genomic sites

We reasoned that the Orc1 IDR could contribute to genomic target-site selection in two different ways. First, the IDR may function as a non-specific avidity element that keeps Orc1 in proximity to chromatin for site-selection by some other mechanism. Alternatively, the IDR may function as the specificity-determining element itself, selecting target sites all on its own. To discriminate between these, we asked whether other chromatin-binding disordered regions could substitute for the human Orc1 IDR. Specifically, we generated human Orc1 chimeras by swapping the native IDR for either the *Drosophila* Orc1 IDR (**Fig. 3A**) or the N-terminal IDR of the uncharacterized *Drosophila* protein Q9VU11 (**Fig. 3B**). These IDRs are relatively similar to the human Orc1 IDR in length (**Fig. 3A-B**) and amino acid composition (**Fig. 3C**), but lack linear sequence similarity, even between the fly and human Orc1 IDR orthologs (*33*).

**Figure 3:**
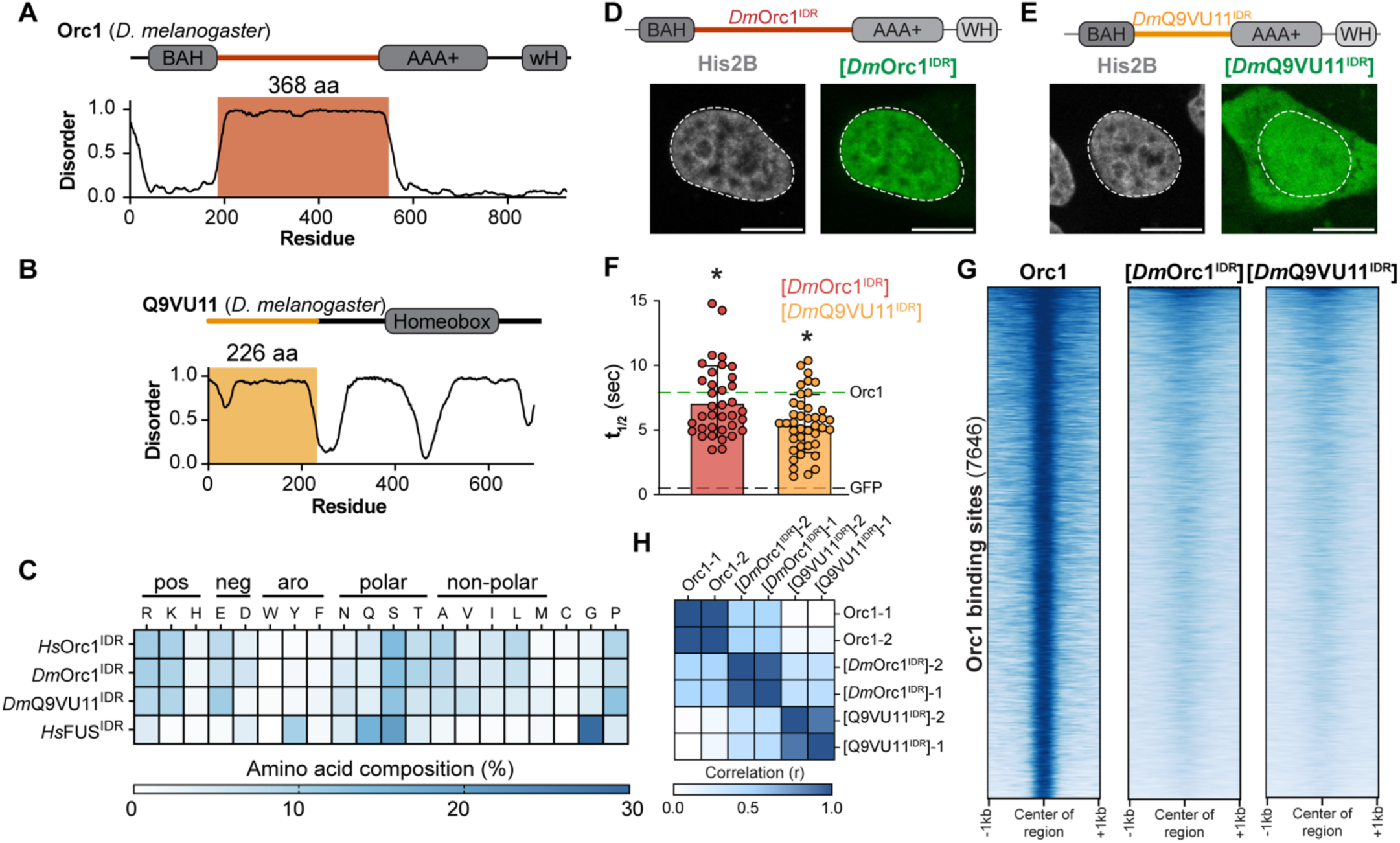
Swapping disordered regions retargets Orc1 to new genomic sites. (A-B) Gene architecture and Metapredict analysis of (A) *Drosophila* Orc1 and (B) *Drosophila* Q9VU11 highlighting their intrinsically disordered regions. (C) Heat map comparing the amino acid composition of the human Orc1 IDR, fly Orc1 IDR (residues 186-549), fly Q9VU11 IDR (residues 2-226), and the N-terminal IDR from human FUS (residues 1-287). (D-E) Representative confocal fluorescence images of (D) Orc1^[*Dm*Orc1-IDR]^ and (E) Orc1^[Q9VU11-IDR]^ chimeric proteins in HeLa cells; scale bar = 10 µm. (F) Half-time of recovery calculated from FRAP analyses of Orc1^[*Dm*Orc1-IDR]^ and Orc1^[Q9VU11-IDR]^ in HeLa cells. Each marker represents a separate cell; the mean and standard deviation are plotted. Significance was established through a one-way ANOVA test compared to GFP (* = p < 0.05). (G) Heat map representation of sequencing read intensity of Orc1^[*Dm*Orc1-IDR]^ (second panel) and Orc1^[Q9VU11-IDR]^ (third panel) across identified Orc1 peaks (first panel, +/− 1 kb). (H) Correlation analysis of CUT&RUN genome coverage files for replicates of Orc1, Orc1^[*Dm*Orc1-IDR]^, and Orc1^[Q9VU11-IDR]^.

We expressed each Orc1 chimera with a C-terminal GFP-Flag tag in HeLa cells to first confirm their nuclear localization. We found that the Orc1 chimera containing the fly Orc1 IDR (Orc1^[*Dm*Orc1-IDR]^) was predominantly nuclear (**Fig. 3D**), while the chimera containing the Q9VU11 IDR (Orc1^[*Q9VU11*-IDR]^) also had a cytoplasmic population (**Fig. 3E**). The chimeras further differed in their nuclear patterning. While Orc1^[*Q9VU11*-IDR]^ appeared to be uniformly distributed in the nucleus, Orc1^[*Dm*Orc1-IDR]^ had the same non-uniform, punctate distribution we and others have observed for human and fly Orc1, which results from its functional enrichment in heterochromatin (*35*, *40*). We next used FRAP to determine whether the chimeric proteins are chromatin-associated and found that both chimeras have a half-time of recovery on par with or slightly faster than wild-type human Orc1 (**Fig. 3F**, Orc1^[*Dm*Orc1-IDR]^ t_1/2_ =7.2 sec ± 2.8 and Orc1^[*Q9VU11*-IDR]^ t_1/2_ = 5.5 sec ± 2.2). These assays confirm that the chimeric Orc1s are enriched in the nucleus and bound to chromatin.

Having confirmed that the swapped IDRs provide an effective substitute in bulk chromatin binding, we next asked how their genome binding specificity compared to wild-type Orc1. Using CUT&RUN, we observed markedly distinct patterns of chromosomal binding between Orc1 and each of the chimeras. Orc1^[*Q9VU11*-IDR]^ was broadly enriched across chromosomes with higher background binding compared to wild-type Orc1 and Orc1^[*Dm*Orc1-IDR]^ (**Fig. S3A**). Further, we observed markedly reduced enrichment of both chimeras at genomic regions corresponding to Orc1 peaks (**Fig. 3G**). Although only weakly present at Orc1 binding sites, both chimeras showed high enrichment at other chromosomal loci, and MACS2 peak calling identified more than 10,000 peaks for Orc1^[*Dm*Orc1-IDR]^ and twice that amount for Orc1^[*Q9VU11*-IDR]^ (**Fig. S3B-C**). Analyzing sequencing reads at specific genomic loci revealed that while some peaks are shared across all three proteins, each chimera also has unique binding sites that are not shared with Orc1 (**Fig. S3A**). Consistent with this, we observed no significant correlation between the genome coverage files for wild-type Orc1 and the two chimeras, although Orc1’s correlation with Orc1^[*Dm*Orc1-IDR]^ (r = 0.60) was substantially higher than with Orc1^[*Q9VU11*-IDR]^ (r = 0.23) (**Fig. 3H**). These data demonstrate that the Orc1 IDR is not functionally interchangeable with other chromatin-binding disordered regions, suggesting a key role for the IDR in guiding site-specific genome binding.

### The Orc1 IDR’s genome targeting activity extends to heterochromatin

ORC is required for the formation and maintenance of heterochromatin (*35*, *41*, *42*), which forms at distinct chromosomal loci and occupies specific sub-nuclear positions (*35*). We therefore asked whether the Orc1 IDR is similarly required for this more cytologically conspicuous form of site-specific chromosome localization. We first confirmed the identity of Orc1 nuclear foci by co-expressing GFP-tagged Orc1 with mCherry-tagged Heterochromatin protein 1 alpha (Hp1*a*) in HeLa (**Fig. 4A**) and HEK293 cells (**Fig. S4A**). In both cell types, we observed strong co-localization of Orc1 with Hp1*a*-enriched regions, both around nucleoli and within peripheral foci (**Fig. 4A**). Similar to the diffuse population of Orc1, FRAP analysis of Orc1’s heterochromatic population revealed that it is dynamically bound to chromatin at these sites (**Fig. 4B**), with a half-time of recovery not appreciably different from the diffusive population of Orc1 (see inset of **Fig. 4B**, t_1/2_ = 8.7 sec ± 2.4).

**Figure 4:**
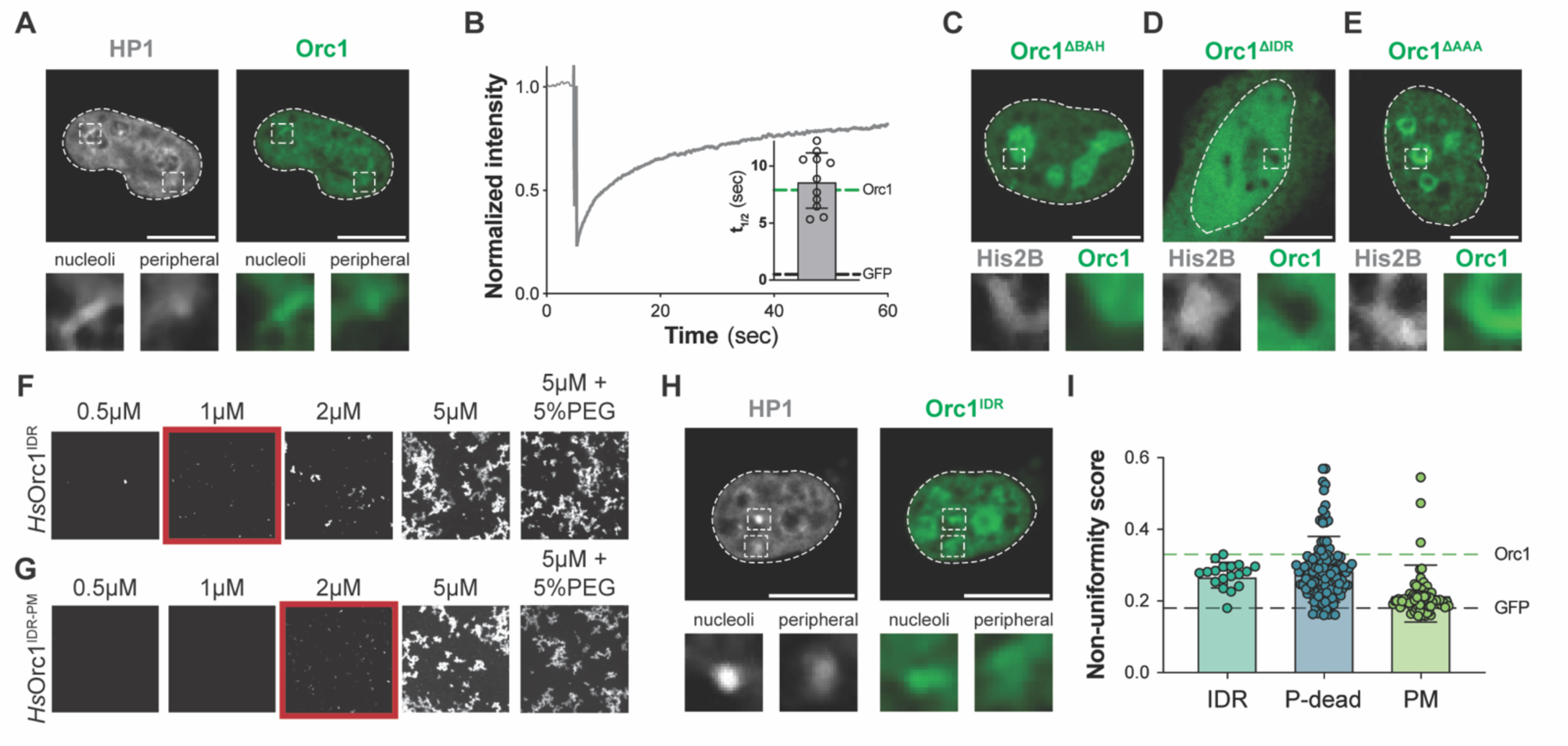
The Orc1 IDR’s genome targeting activity extends to heterochromatin. (A) Confocal fluorescence imaging showing nuclear distribution of mCherry-tagged Heterochromatin protein 1*a* (Hp1*a*) and GFP-tagged human Orc1 in HeLa cells. Zoom windows include nucleolus-proximal foci (region I) and peripheral foci (region II). Scale bar = 10 µm. (B) Analysis of GFP-Orc1 FRAP dynamics in foci. Inset shows the calculated half-time of recovery compared to the non-foci population (green line, from Fig. 1C). (C-E) Confocal fluorescence images of Orc1 domain deletion constructs, including (C) Orc1^ΔBAH^, (D) Orc1^ΔIDR^, and (E) Orc1^ΔAAA^. Zoom windows show regions of heterochromatin as indicated by the high-intensity mCherry-H2B signal. Scale bar = 10 µm. (F-G) Phase separation screen of (F) Orc1^IDR^ and (G) Orc1^IDR-PM^ in the presence of stoichiometric amounts of Cy5-labeled 60 bp duplex DNA. Phase separation was also tested in the presence of PEG. (H) Confocal fluorescence imaging showing the nuclear distribution of mCherry-Hp1*a* and GFP-Orc1^IDR^ in HeLa cells. Zoom windows show nucleolar-proxomal foci (region I) and peripheral foci (region II). (I) Comparison of nuclear non-uniformity (n.u.) between Orc1^IDR^, Orc1^IDR-P-dead^, and Orc1^IDR-PM^. Significance established with one-way ANOVA compared to Orc1^IDR^ (* = p < 0.05).

We next asked which domain(s) of Orc1 is required for heterochromatin localization. Specifically, we expressed GFP-tagged Orc1 domain deletion constructs (Orc1^ΔBAH^, Orc1^ΔIDR^, and Orc1^ΔAAA^) in HeLa cells and, in addition to visually assessing heterochromatin localization (**Fig. 4C-E**), we quantified the level of foci formation by calculating the non-uniformity of the fluorescence signal in the nucleus (**Fig. S4B**). We found that removal of Orc1’s folded domains did not appear to affect Orc1 recruitment to heterochromatin (**Fig. 4C, E**), although removal of the BAH domain did drive partitioning of Orc1^ΔBAH^ into nucleoli. In contrast, removal of the IDR completely abolished foci formation (**Fig. 4D**) and image quantification showed that while full-length Orc1, Orc1^ΔBAH^, and Orc1^ΔAAA^ had similarly high non-uniformity scores (Orc1 n.u. = 0.33 ± 0.10, Orc1^ΔBAH^ n.u. = 0.32 ± 0.07, Orc1^ΔAAA^ n.u. = 0.30 ± 0.08), the score for Orc1^ΔIDR^ was significantly reduced and similar to the negative control (Orc1^ΔIDR^ n.u. = 0.18 ± 0.02, GFP n.u. = 0.21 ± 0.13, **Fig. S4B**). These results demonstrate that the Orc1 IDR is necessary for heterochromatin targeting.

Heterochromatin is believed to form through phase separation of its resident factors (*43*, *44*), and prior work shows that metazoan ORC can phase separate in the presence of DNA (*45*, *46*). In flies, ORC’s phase separation depends on the Orc1 IDR. We therefore asked whether the human Orc1 IDR is sufficient to drive phase separation and heterochromatin partitioning. Interestingly, we found that the human Orc1^IDR^, when combined with stoichiometric amounts of DNA, formed a protein network rather than a condensed liquid phase (**Fig. 4F**). We observed the same type of assemblies with a purified phospho-mimetic Orc1 IDR (Orc1^IDR-PM^), although it had a significantly elevated critical concentration (**Fig. 4G**). The impact of phosphorylation is consistent with prior work showing a phosphorylation-dependent reduction in fly ORC’s phase separation propensity (*40*, *45*). We next asked whether the IDR, being sufficient for self-assembly *in vitro*, is likewise sufficient for heterochromatin partitioning. Similar to full-length Orc1, we found that Orc1^IDR^ co-localized with HP1*a* around nucleoli and within other subnuclear foci (**Fig. 4H**), although its non-uniformity (Orc1^IDR^ n.u. = 0.27 ± 0.03, **Fig. 4I**) was slightly lower than what was observed for the full-length protein. We also assessed the impact of phosphorylation on heterochromatin partitioning and found that an Orc1 IDR variant that cannot be phosphorylated (Orc1^IDR-P-dead^) had a small increase in non-uniformity compared to Orc1^IDR^ (Orc1^P-dead^ n.u. = 0.29 ± 0.09), while a phospho-mimetic showed a strong reduction in non-uniformity (Orc1^PM^ n.u. = 0.22 ± 0.08) (**Fig. 4I**), behaving more similarly to GFP than to Orc1. A similar effect was observed in HEK293 cells, with some nuanced differences. In this cell type, heterochromatic foci were more abundant, the IDR showed higher levels of non-uniformity, and phosphorylation exerted a more dominant effect (**Fig. S4C-F**). Taken together, these results demonstrate that the Orc1 IDR is necessary and sufficient for heterochromatin targeting and that this activity is negatively regulated by phosphorylation.

### Licensing factor IDRs have non-equivalent roles in genome targeting

Several other replication licensing factors have long IDRs, raising the question of whether the Orc1 IDR is unique or if licensing factor IDRs generally possess chromatin binding and genome targeting activity. Specifically, within the human ORC complex, Orc1 (IDR = 319 aa), Orc2 (IDR = 243 aa) and Orc6 (IDR = 67 aa) each have a disordered region longer than 50 amino acids, as do Pre-RC components Cdt1 (IDR = 175 aa) and Cdc6 (IDR = 142 aa) (**Fig. 5A**). Licensing factor IDRs are of relatively high sequence complexity (**Fig. S5A**) (*33*) with nearly all amino acid types represented at appreciable levels (except aromatic residues, **Fig. 5B**). As a result, it is difficult to meaningfully compare the IDRs’ amino acid compositions to determine if they are more or less similar to the Orc1 IDR. However, Pre-RC IDRs can be broadly grouped by their charged residue content: Orc1, Cdt1, and Cdc6 IDRs bear a net positive charge (between +12.1 and +29.5) with charged residues equitably distributed throughout the sequence (**Fig. S5B**), while Orc2 and Orc6 have a weak net negative charge (between −4.7 and −2.5) (**Fig. 5C**). Given the high net positive charge of the Orc1 IDR, which is consistent with its DNA and chromatin binding activity, we tested whether the Cdt1 and Cdc6 IDRs may likewise have a role in chromatin binding and genome targeting.

**Figure 5:**
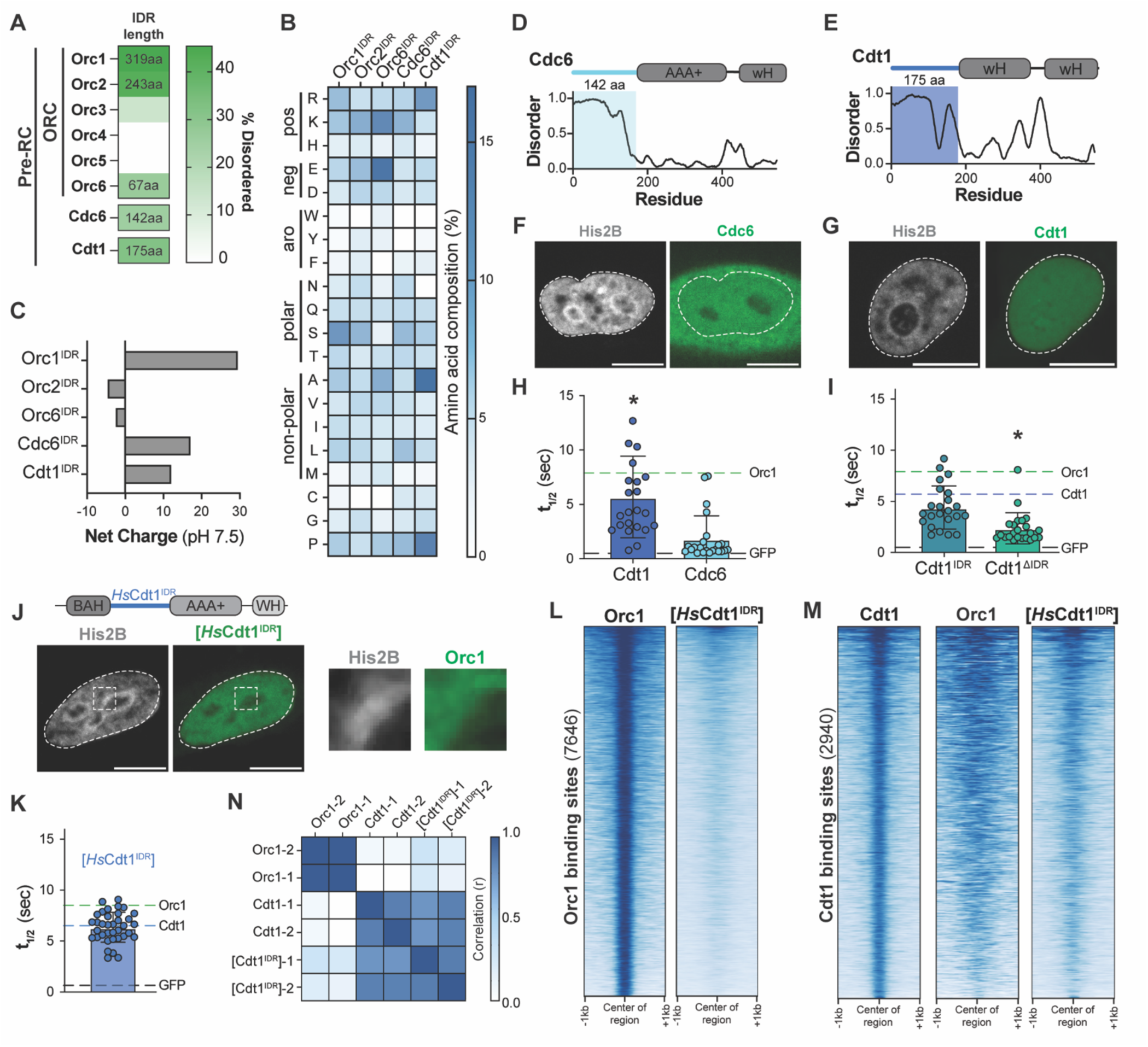
Licensing factor IDRs have non-equivalent roles in genome targeting. (A) Analysis of licensing factor disorder content, include total percent disordered (green) and longest disordered region. (B) Heat map comparing the amino acid composition of the human Orc1 IDR, Orc2 IDR, Orc6 IDR, Cdc6 IDR, and Cdt1 IDR. (C) Net charge of Orc1, Orc2, Orc6, Cdc6, and Cdt1 disordered regions. (D-E) Domain architecture and Metapredict disorder prediction highlighting the IDRs of (D) Cdc6 and (E) Cdt1. (F-G) Cellular distribution of GFP-tagged (F) Cdc6 and (G) Cdt1 in HeLa cells stably expression mCherry-H2B; scale bar = 10 µm. (H) Half-time of recovery calculated from FRAP experiments with GFP-tagged Cdt1 and Cdc6. Significance calculated using a one-way ANOVA and compared to GFP (black dotted line) (* = p < 0.05). (I) Half-time of recovery for Cdt1^IDR^ and Cdt1^ΔIDR^. Significance calculated using a one-way ANOVA and compared to full-length Cdt1 (blue dotted line) (* = p < 0.05). (J) Domain architecture of Orc1^[*Hs*Cdt1-IDR]^ chimera and its nuclear distribution in HeLa cells stably expressing mCherry-tagged H2B. Zoom window shows nucleoli-proximal H2B foci. Scale bar = 10 µm. (K) Half-time of recovery for Orc1^[*Hs*Cdt1-IDR]^. Significance calculated using a one-way ANOVA and compared to Cdt1 full-length (blue dotted line) (* = p < 0.05). (L) Heatmap representation of sequencing read intensity of Orc1^[*Hs*Cdt1-IDR]^ (second panel) across Orc1 peaks (first panel, +/− 1 kb). (M) Heatmap representation of sequencing read intensity of Orc1 (second panel) and Orc1^[*Hs*Cdt1-IDR]^ (third panel) across Cdt1 peaks (first panel, +/− 1 kb). (N) Correlation analysis of CUT&RUN genome coverage files for replicates of Orc1, Cdt1, and Orc1^[*Hs*Cdt1-IDR]^.

We expressed Cdc6 (**Fig. 5D**) and Cdt1 (**Fig. 5E**) with a C-terminal GFP-Flag tag and assessed their cellular localization (**Fig. 5F-G**) and chromatin binding dynamics (**Fig. 5H**). Consistent with prior reports (*34*, *47–49*), we found that Cdc6 was present in both the cytosol and nucleus (**Fig. 5F**) while Cdt1 was exclusively nuclear (**Fig. 5G**). Interestingly, neither appeared to be enriched within heterochromatin and only Cdt1 showed evidence of chromatin association as assessed by FRAP (**Fig. 5H**, Cdt1 t_1/2_ = 5.7 ± 3.7 sec, Cdc6 t_1/2_ = 1.8 ± 2.1 sec) despite both having IDRs of relatively equal length and the IDR of Cdc6 bearing a higher net positive charge (**Fig. 5C**). To investigate the contribution of the Cdt1 IDR to chromatin association, we quantified the chromatin binding dynamics of either the N-terminal IDR alone (Cdt1^IDR^) or the C-terminal folded domains. Notably, deletion of the IDR abolished Cdt1’s nuclear enrichment (**Fig. S5C**), and therefore a nuclear localization signal was added to our IDR deletion construct (Cdt1^NLS-ΔIDR^). Both constructs showed faster recovery rates compared to full-length Cdt1 (**Fig. 5I**, Cdt1^IDR^ t_1/2_ = 4.4 ± 2.1 sec, Cdt1^NLS-ΔIDR^ t_1/2_ = 2.4 ± 1.5 sec), consistent with weakened chromatin association. These findings agree with prior *in vitro* and live-cell quantitative microscopy studies that demonstrate a contribution from the Cdt1 N-terminus and first winged-helix domain to its DNA and chromatin binding activity (*34*, *50*). Like our observations with Orc1 (**Fig. 1-2**), this suggests that Cdt1’s folded domains and IDR cooperate to achieve maximal chromatin engagement.

We next investigated the contribution of the Cdt1 IDR to genomic targeting using the same chimera-based approach we used previously. We generated an Orc1 chimera containing the Cdt1 IDR (Orc1^[HsCdt1-IDR]^, **Fig. 5J**) and assessed chromatin binding and genome localization. It was immediately apparent from cellular studies that Orc1^[HsCdt1-IDR]^ no longer behaved like wild-type Orc1 but had become more Cdt1-like. Specifically, it was no longer enriched within heterochromatin (**Fig. 5J**) and possessed chromatin binding dynamics (**Fig. 5K**, Orc1^[HsCdt1-IDR]^ t_1/2_ = 6.4 ± 1.5 sec) more similar to Cdt1 (Cdt1 t_1/2_ = 5.7 sec) than Orc1 (Orc1 t_1/2_ = 7.9 sec). CUT&RUN analysis of the chimera revealed a loss of Orc1^[HsCdt1-IDR]^ signal at full-length Orc1 peaks (**Fig. 5L**), confirming the importance of the native Orc1 IDR in guiding target site selection. To determine whether it is Cdt1’s disordered region that guides genome targeting, we used CUT&RUN to map Cdt1’s genomic binding sites and assessed wild-type Orc1 and Orc1^[HsCdt1-IDR]^ signal over these regions. Although we observed Orc1 occupancy within a 2 kb region surrounding Cdt1 peaks, the Orc1 signal was disorganized and not centered on Cdt1 binding sites (**Fig. 5M**, compare left and middle panels). Consistently, there was no correlation between the Cdt1 and Orc1 genome coverage files (**Fig. 5N**). However, we found that the Orc1^[HsCdt1-IDR]^ chimera displayed strong, focused enrichment centered around wild-type Cdt1 binding sites (**Fig. 5M**, compare left and right panels), and Pearson correlation analysis confirmed that the genomic binding pattern of the chimera was highly correlated with Cdt1 (**Fig. 5N**, r = 0.8). These data demonstrate that the Cdt1 IDR, like the Orc1 IDR, directs chromatin binding specificity and that this function does not depend on the other folded chromatin-binding domains in the protein.

## DISCUSSION

Here we demonstrate a key role for the Orc1 disordered region in guiding genome binding specificity. We find that removal of the IDR abolishes chromatin association and that the IDR’s function extends beyond non-specific DNA binding to guide Orc1 to specific genomic loci. This function is dependent on the identity of the IDR: swapping the human Orc1 IDR for other chromatin-binding disordered regions redirects the protein to new genomic sites. We additionally show that the Orc1 IDR is necessary and sufficient for heterochromatin recruitment, demonstrating its utility across heterogeneous chromatin architectures. While the IDR plays the leading role in specificity, Orc1’s folded domains, and especially the histone binding BAH domain, are nonetheless essential and appear to function synergistically with the IDR to increase the protein’s retention time on chromatin. Finally, we show that DNA replication licensing factor IDRs have non-equivalent functionalities, with the IDR of Cdt1 but not Cdc6 able to bind chromatin and direct genomic localization, as evidenced by the redirection of an Orc1 chimera containing the Cdt1 IDR to Cdt1 binding sites. Together, our findings support a model in which licensing factor IDRs function as information-rich chromatin-binding elements that direct site-specific chromatin binding.

Origin selection mechanisms have evolved remarkably across eukaryotes and have been most thoroughly studied in budding and fission yeast. In both yeast species, origins are defined in cis, but their origin recognition mechanisms differ. In budding yeast, origin selection is largely dictated by a species-specific helical insertion in Orc4 that recognizes an 11-base pair DNA motif, the ARS consensus sequence (ACS) (*22*, *51*). In fission yeast, origin specification shifts away from a specific motif towards more general DNA sequence features with compensatory changes to ORC’s architecture: the *S. pombe* Orc4 ortholog has a long N-terminal disordered region containing multiple AT-hook motifs that preferentially bind the minor groove of asymmetric AT-rich DNA (*24*, *52*). Studies of metazoan ORC suggest a more complex mechanism. For example, while metazoan ORC retains the ability to bind specific genomic loci, these sites do not share a common DNA sequence motif (*26*, *27*, *53*) and, consistently, metazoan ORC binds DNA promiscuously (*25*, *54*). An additional point of distinction is the labile nature of the human ORC holocomplex, which exists as sub-assemblies within the nucleus (Orc1, Orc2-5, and Orc6). As a result of these complexities, the mechanism underlying origin recognition by metazoan ORC has long remained a mystery.

Each of human ORC’s dissociable complexes could, in theory, bind and nucleate Pre-RC assembly at specific genomic loci. In fact, Orc1, Orc2-5, and Orc6 can each independently bind DNA and/or chromatin (*30*, *31*, *45*, *55*). While subcomplex-specific origin specification warrants investigation, multiple lines of evidence point to an exceptional role for Orc1. First, Orc1 contains both a histone-binding BAH domain (*31*) and a DNA-binding IDR (*45*). The importance of these elements is emphasized by known disease-causing mutations in the human BAH domain (*38*) and, for the IDR, its requirement for chromatin recruitment and viability in flies (*33*, *56*). Further, prior work demonstrates that chromatin recruitment of human Orc2-5 depends on the presence of Orc1 (*29*, *37*). We therefore focused our efforts on understanding how Orc1’s multiple chromatin binding elements coordinate site-specific chromatin engagement. As opposed to flies, where the BAH domain is dispensable (*45*), we found that the human Orc1 IDR and BAH domain function synergistically, with deletion of either element reducing bulk chromatin association. The species-specific dependence on the Orc1 BAH domain highlights the evolutionary plasticity of metazoan origin selection mechanisms and is in line with recent work that demonstrates mechanistic divergence in downstream licensing steps (*57*). Conversely, deletion of the AAA+ domain had no measurable effect on Orc1’s FRAP recovery dynamics, suggesting that formation of and DNA encirclement by the ORC holocomplex likely occurs downstream of origin selection and may be a transient state. In terms of genome binding specificity, we found that the IDR, not the BAH domain, plays the leading role. Indeed, Orc1’s genome binding specificity was retained in the absence of the BAH domain (**Fig. 1**) but not the IDR (**Fig. 2**), and Orc1 chimeras that left the BAH domain intact but swapped IDRs were redirected to new genomic locations (**Fig. 3** and **Fig. 5**).

Based on these results, we propose a three-step model for human origin licensing (**Fig. 6**): 1) **Scanning** – we and others have demonstrated that Orc1 is dynamically associated with chromatin (*35*). These dynamic interactions are driven by synergy between the BAH domain and IDR, and we predict that this enables Orc1 to rapidly scan the genome for specific binding sites. Owing to the abundance of the epigenetic mark to which the BAH domain binds (H4K20me2, present on >80% of nucleosomes genome-wide (*58*)), and the non-specific DNA-binding activity of the IDR (*33*), we predict that Orc1 can interact with nearly any region of chromatin in this scanning mode. 2) **Recognition** – our data demonstrate that the Orc1 IDR guides site-specific chromatin binding. These sites are not defined in cis (*26*, *53*), which is consistent with the IDR’s promiscuous nucleic acid-binding activity, and we thus predict that specificity is determined by a hitherto unidentified trans-acting factor(s) (discussed below). Orc1’s dynamic association with chromatin suggests that this trans interaction is likely weak and might only marginally extend the lifetime of Orc1 on chromatin to increase the probability of a licensing-competent engagement at specific chromosomal loci. 3) **Capture and assembly** – the lifetime of Orc1 on chromatin is likely a key determinant of licensing efficiency since multiple, possibly rare conditions must be met for a functional association with the genome: a) Orc1 must capture the remaining ORC subunits (Orc2-5 and Orc6); b) the ORC holocomplex must encircle DNA, stipulating the presence of approximately 40 base pairs of naked DNA nearby (*59*); and c) the remaining Pre-RC components (Cdc6, Cdt1, and Mcm2-7) must bind and assemble on an even longer stretch of unoccupied DNA (> 100 basepairs (*60*)). Notably, the average linker DNA length between adjacent nucleosomes is relatively short (≅40 base pairs (*61*)) and, without a dedicated histone chaperone, ORC and Pre-RC assembly may have to rely on serendipitous chromatin opening (*26*, *62*), emphasizing the importance of Orc1’s binding kinetics.

**Figure 6:**
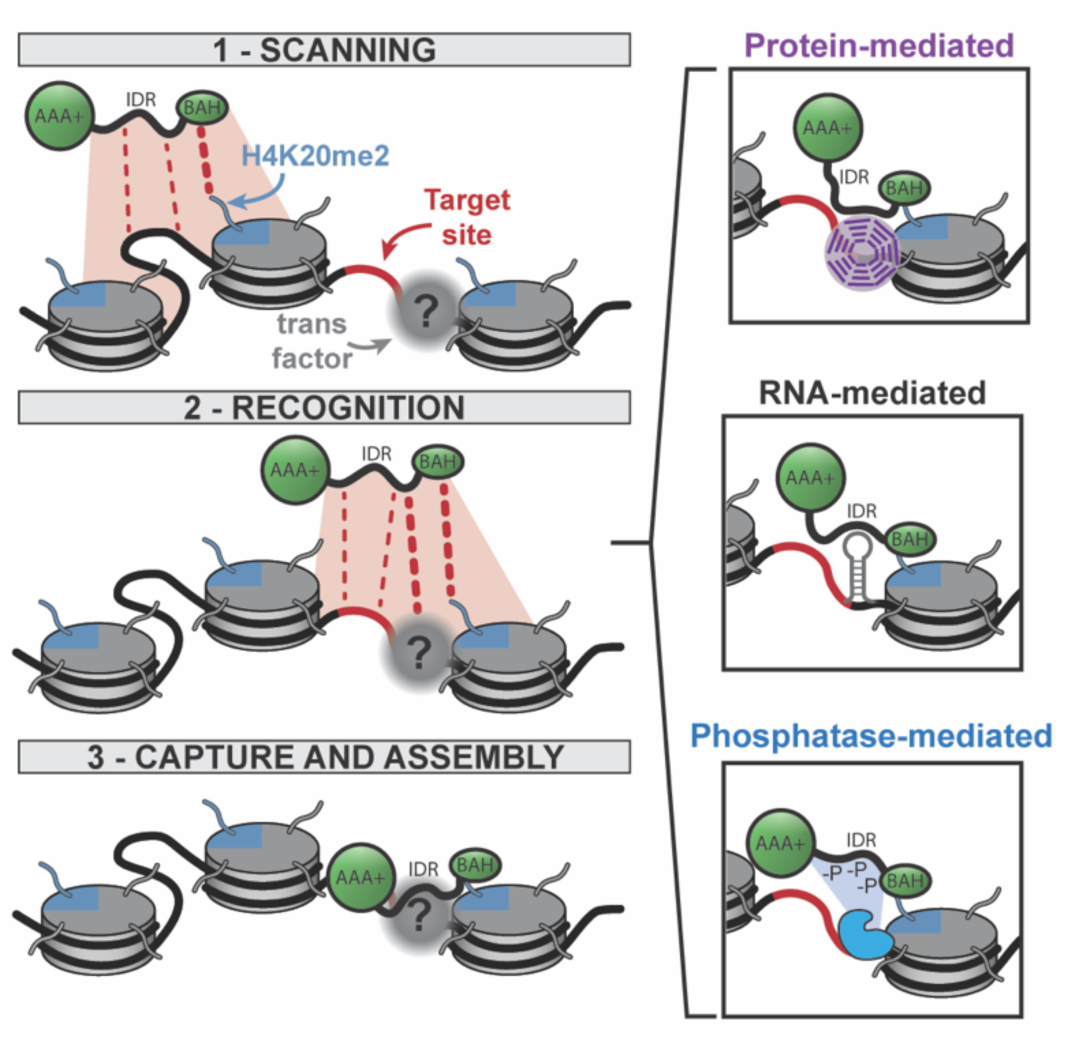
Model for the functional association of Orc1 with chromatin. Orc1 identifies genomic binding sites in three steps: 1) genome scanning facilitated by the synergistic chromatin-binding activities of the Orc1 BAH domain and IDR, 2) recognition of specific genomic loci by IDR-mediated interactions with a chromatin-associated factor(s), and 3) functional chromatin association by capture of the remaining ORC subunits and full Pre-RC assembly.

The mechanism of IDR-mediated origin recognition remains an open question, including the nature and identity of any trans-acting factor(s). We see three non-mutually exclusive possible mechanisms (**Fig. 6**). First, Orc1 may recognize chromatin-associated proteins via short linear motifs (SLiMs) contained within the Orc1 IDR. Known Orc1 IDR SLiMs include binding sites for Cyclin-Dependent Kinases (CDK) and Protein Phosphatase 1 (PP1) (*46*). These known SLiMs account for only a small fraction of the Orc1 IDR length, and the remaining sequence, which is of unknown significance, may encode additional protein-protein interaction motifs. In support of this, SLiMs were recently identified in the fly Orc1 IDR that mediate binding to Hp1 (*42*), and this may account for the IDR’s heterochromatin-targeting activity (**Fig. 4**). Second, the Orc1 IDR may recognize specific nucleic acid species. Although the IDR binds duplex DNA non-specifically (*33*), RNAs are also enriched on chromatin (*63*), and their structural heterogeneity may favor unrecognized degrees of specificity in IDR-nucleic acid interactions. In fact, this mode of origin recognition is operative in at least three specific cases: at the Epstein-Barr Virus (EBV) origin of replication (OriP) (*64*), at telomeres (*65*), and at the *c-MYC* replication origin (*66*). In each case, a specific RNA facilitates recruitment of Orc1. Third, specificity could be conferred through localized regulation of the Orc1 IDR’s non-specific DNA-binding activity, which is inhibited by CDK-dependent phosphorylation (*33*). We have recently shown that phosphorylation of the fly Orc1 IDR inhibits its recruitment to heterochromatin (*40*), a mechanism we find to be conserved in human cells. Notably, Orc1’s regulatory phosphatase, PP1, is locally enriched within heterochromatin due to its interaction with Rap1-interacting factor 1 (Rif1) (*67–69*). This led us to hypothesize that PP1, which is tethered to chromatin through its interaction with various adaptor proteins (*70–73*), might locally dephosphorylate Orc1 to unveil its non-specific DNA-binding activity in specific genomic locations. This may explain why the Orc1 chimera containing the fly Oc1 IDR can partially rescue the genome binding specificity of the wild-type protein, as the fly Orc1 IDR also contains a PP1 docking motif (*40*). Altogether, these observations point to the Orc1 IDR functioning as an integrative interaction scaffold that senses local chromatin cues to guide site-specific genome binding in the absence of a traditional sequence-specific DNA binding domain.

These findings add to a growing body of work emphasizing the importance of intrinsically disordered regions in genome navigation (*15*, *17*, *33*, *45*, *74*). This functional role of IDRs was first appreciated in transcription factors (*15*), and our observations now extend this to the process of DNA replication licensing. These findings raise the broader question of the generality of IDR-mediated genome targeting in nuclear proteins. Notably, long disordered regions are overrepresented in the nuclear proteome (*75*, *76*) and replication licensing factors are no exception: three ORC subunits as well as Cdt1 and Cdc6 have relatively long IDRs (length range = 67-319 amino acids, **Fig. 5**). When we compared the function of Orc1, Cdc6, and Cdt1 IDRs, we found that only the Cdt1 and Orc1 IDR could facilitate chromatin binding and direct genome localization. Although limited in scope, these findings suggest that the disordered regions of nuclear proteins likely have highly individualized functions, even for IDRs within the same pathway and with similar sequence features (e.g., sequence composition, net charge, and sequence complexity). Indeed, our data show that while both the Orc1 and Cdt1 IDRs can bind chromatin, they encode unique specificities. Beyond IDR primary structure, functional distinctions may also depend on the specific protein context in which an IDR is found. For example, it is interesting that a distinguishing feature of Orc1 and Cdt1 is the presence of a general DNA/chromatin binding element: the Orc1 BAH domain, which binds an abundant epigenetic mark, and Cdt1’s winged-helix domains, which bind DNA directly (*50*). The chromatin interactions contributed by these domains may critically support the function of the IDR.

In summary, we have demonstrated that human Orc1 recognizes genomic binding sites via its intrinsically disordered region. This work provides fresh insight into the mechanism of sequence-independent origin specification in human cells and paves the way for future work to uncover the chromatin-associated factors recognized by the Orc1 IDR. Together with studies of transcription factor site selection, this work reveals that IDR-mediated genome navigation is important for multiple genomic processes, and an important future goal is to understand the prevalence of this genome targeting strategy and the mechanisms involved. Clearly, protein disordered regions possess an impressive level of functional sophistication, and studies of their genome targeting activity point to highly specific and intricate modes of molecular recognition. These mechanisms seem likely to depend on the ability of IDRs to integrate diverse chromatin cues and discerning the hierarchy of interactions that guide IDR genome targeting is an exciting future frontier.

## MATERIALS AND METHODS

### Molecular cloning

Licensing factor imaging constructs were prepared by cloning each factor and subdomain construct into the SspI restriction site of vector 6D (QB3 Macrolab) using ligation-independent cloning. This vector was custom-modified via around-the-horn mutagenesis to introduce a C-terminal FLAG tag (DYKDDDDK). Specifically, we cloned full-length human Orc1, Cdt1, and Cdc6, in addition to the following sub-domains and domain deletion constructs: Orc1^ΔBAH^ (deletion of residues 1-176), Orc1^ΔIDR^ (deletion of residues 177-484), Orc1^ΔAAA^ (deletion of residues 485-861), Orc1^BAH^ (residues 1-176), Orc1^IDR^ (residues 177-484), Orc1^AAA^ (residues 485-861), Cdt1^ΔIDR^ (deletion of residues 1-165), Cdt1^NLS-ΔIDR^ (N-terminal SV-40 NLS: PKKKRKV), and Cdt1^IDR^ (residues 1-166). Orc1 chimeras were generated by Gibson Assembly (NEB, Cat. #E2611S) reactions that swapped the coding sequence for the human Orc1 IDR (residues 177-484) with the sequence of the *Drosophila* Orc1 IDR (residues 186-549), the Q9VU11 IDR (residues 2-226), or the human Cdt1 IDR (residues 2-166). The Hp1*a* coding sequence was cloned into vector 6B (QB3 MacroLab) for expression as a C-terminal mCherry fusion using the same LIC strategy. For generating purified proteins, the coding region of the human Orc1 IDR (residues 177-484) was cloned into vector 4C (Macrolab) for expression in Sf9 cells as a TEV-cleavable His6-MBP fusion protein. The Orc1^IDR-P-dead^ construct was generated by synthesizing (Twist Biosciences) a construct with every “[S/T]P” motif changed to “AP” (residues: S199A, T203A, T224A, T230A, S258A, S273A, S287A, S311A, T337A, T375A, T428A, T466A). Similarly, Orc1^IDR-PM^ was synthesized with the same serine or threonine residues mutated to aspartate. Both constructs were cloned into vector 4C for expression and purification or into vector 6D for imaging studies. All constructs were verified by whole-plasmid sequencing (Plasmidaurus).

### Cell Culture

HeLa Kyoto cells and the same stably expressing Histone H2B-mCherry (gifts from the Laboratory of Dr. Michael Rosen) were maintained in Dulbecco’s Modified Eagle Medium (DMEM; Gibco, Cat. #11965-092) supplemented with 10% fetal bovine serum (FBS; Gibco, Cat. #A52568-01) and 1% Penicillin–Streptomycin–Glutamine (Gibco, Cat. #10378-016). Cells were cultured at 37°C in a humidified incubator with 5% CO₂. Cells were passaged into a µ-Plate 24-well ibiTreat (Ibidi, Cat. #82426) glass bottom imaging dish at a 1:10 ratio one day before transfection. Plasmid transfections were performed using jetPRIME transfection reagent (Sartorius, Cat. #101000015) according to the manufacturer’s instructions. Briefly, for each well of a 24-well plate, 0.5 μg of plasmid DNA was mixed with 50 μL of jetPRIME buffer and 1 μL of jetPRIME reagent. The mixture was incubated for 15 minutes at room temperature before being added dropwise to cells. Live-cell imaging was performed 24 hours post-transfection.

### Cell fractionation and immunoblotting

24 hours post transfection cells were collected and either used to prepare a whole-cell lysate or fractionated. For whole-cell lysates, cells were washed with 1 mL PBS and centrifuged at 600g for 3 minutes. Next, cells were resuspended in 30 µL of RIPA buffer (25mM Tris, 150mM NaCl, 1% NP40, 1% Deoxycholate, 0.1% SDS) and incubated on ice for 30 min. Finally, cells were spun down at 14,000 RPM for 15 min. For fractionation, cells were washed with PBS as described above and then fractionated using a standard kit following the manufacturer’s instructions (Abcam, Cat. #ab219277). Briefly, cells were resuspended in cytoplasm isolation buffer, centrifuged at 1000 g for 3 minutes, followed by resuspension in nuclear soluble buffer and centrifugation at 5000 g for 3 minutes, and finally resuspended in nuclear bound buffer. The supernatant was collected for both fractionated and whole-cell lysate samples and mixed with 2x Laemmli loading buffer in a 1:1 v/v ratio. Samples were heated to 95°C and loaded onto a mini-PROTEAN stain-free protein gel (Bio-Rad, Cat. #4568096). Electrophoresis was run at 300V for 17 min, followed by transfer using the Trans-Blot Turbo Transfer system (Bio-Rad, Cat. #1704158) at 1.3 A, 25 V for 7 min. Membranes were washed in TBS-T buffer three times for 2 min each, then blocked with 5% BSA (GoldBio, Cat. #A-420-500) for 30 min. Next, membranes were washed three times with 1x TBS-T buffer for 5 min each, and incubated with primary antibody in TBS-T at 4°C overnight. The next day, the membranes were washed three times for 5 min each with TBS-T and then incubated with secondary antibody in 5% milk for 30 min at room temperature with shaking, covered in foil. Finally, the membranes were washed three times with 1x TBS-T for 15 min each and then imaged using a ChemiDoc MP Imaging System (Bio-Rad).

### Imaging by confocal fluorescence microscopy

Live-cell imaging was performed on a Nikon Ti2E microscope equipped with a Yokogawa CSU X1 spinning disk using a 60× oil immersion objective (NA=1.4) with the appropriate filter set and an environmental chamber maintained at 37°C and 5% CO₂. Images were acquired using the 488 nm and 561 nm laser lines at 10% laser power with an exposure time of 100 ms. Images were collected with a field of view of 1024 × 1024 pixels. For each construct, a 3 × 3 tiled image set with 10% overlap between adjacent fields was acquired and automatically stitched to generate a composite image. Images were processed using FIJI. Unless otherwise indicated, identical acquisition settings were used for all samples within an experiment. For foci visual assessment, background was subtracted from the cropped image, and a median filter was applied at 2 pixels.

### Fluorescence recovery after photobleaching (FRAP) experiments

FRAP analyses were performed on at least 9 cells per construct. A 1.8 µm x 1.8 µm region of interest (ROI) within the nucleus was photobleached using a 405 nm laser at 25% laser power and a 100 μs dwell time. A five-second delay was introduced before bleaching to acquire pre-bleach images. Image acquisition continued for 60 seconds, yielding 600 frames. Images were acquired at a resolution of 512 × 512 pixels. Fluorescence recovery curves were analyzed using Excel. Intensities were corrected for background fluorescence and photobleaching and normalized to the first frame (t_0_) pre-bleach intensity. Data were analyzed using GraphPad Prism 11. Statistical significance was determined using one-way ANOVA, as appropriate. Error bars represent standard deviation. Statistical significance was defined as p < 0.05, unless otherwise indicated.

### Calculation of nuclear non-uniformity scores

Foci formation was quantified using a custom macro in Fiji. The H2B-mCherry signal was used to generate nuclear masks, and the mean and maximum GFP fluorescence intensities were measured within each nucleus. Cells with a mean GFP intensity below 125 (non-expressing cells) or above 4095 (highly overexpressing cells) were excluded from further analysis. The measured intensities were used to calculate a non-uniformity score (*77*): non-uniformity (n.u.) = (maximum intensity − mean intensity) / (maximum intensity + mean intensity). Data were analyzed using GraphPad Prism 11. Statistical significance was determined using one-way ANOVA, as appropriate. Error bars represent standard deviation. Statistical significance was defined as p < 0.05, unless otherwise indicated.

### Protein purification

The human Orc1 IDR (Orc1^IDR^) and phospho-mimetic (Orc1^IDR-PM^) and phospho-defective (Orc1^IDR-P-dead^) variants were expressed and purified from Sf9 cells. Briefly, bacmid DNA was generated according to established procedures (*45*) and subsequently transfected into Sf9 cells (Expression Systems) maintained in ESF921 (Expression Systems, Cat. #96-001-01) using Cellfectin II (Gibco, Cat. #16362100) following the manufacturer’s instructions. P0 virus was harvested from the transfected cells and amplified twice (to P2 virus) before infection of Sf9 cells for protein expression. Cells were infected and harvested after 2 days. Cell pellets were isolated by centrifugation (15 min, 5000 RPM) and frozen at −80°C until protein purification.

Orc1^IDR^, Orc1^IDR-PM^ and Orc1^IDR-P-dead^ were purified from 2 L of cultured cells. The cell pellet was resuspended in 100 mL of lysis buffer (50 mM Tris, pH 7.5, 300 mM KCl, 50 mM imidazole, 10% glycerol, 200 μM phenylmethylsulfonyl fluoride (PMSF), 1 mM β-mercaptoethanol (BME), 1 μM benzonase (Milipore, Cat. #70664-10KUN) and 1× cOmplete EDTA-free Protease Inhibitor Cocktail (Thermoscientific, Cat. #A32965)). Cells were lysed by sonicating on ice (five cycles of 15 sec at 100% power followed by 1 min rest), and the lysate was centrifuged at 30,000 xg for 1 hr at 4°C. The supernatant was collected and filtered using an aPES 0.45 μm bottle-top filter unit (Nalgene Rapid-Flow, Thermo Fisher, Cat. #FB12566511). The filtrate was loaded onto a 5 ml HisTrap HP column (GE Healthcare, Cat. #17524802), washed with 10 column volumes (CV) of wash buffer (50 mM Tris, pH 7.5, 300 mM KCl, 50 mM imidazole, 10% glycerol, 1 mM BME) and the protein eluted with a 6 CV linear gradient from 0% to 100% elution buffer (50 mM Tris, pH 7.5, 300 mM KCl, 250 mM imidazole, 10% glycerol, 1 mM BME). Amylose purification was used as a second affinity purification step. The sample was loaded onto a column packed with amylose resin (NEB, Cat. #E8021) and subsequently washed with 3 CV of amylose wash buffer (50 mM Tris, pH 7.5, 300 mM KCl, 10% glycerol, 1 mM BME) and eluted with 2 CV of amylose elution buffer (50 mM Tris, pH 7.5, 300 mM KCl, 10% glycerol, 1 mM BME, 20 mM maltose). Finally, the protein was concentrated to < 2 ml using an Amicon Ultra-15 concentrator (Millipore, Cat. #UFC903008) and then purified by size-exclusion chromatography. Samples were loaded onto an Enrich 650 and run with 1.2 CV of sizing buffer (50 mM HEPES, pH 7.5, 300 mM potassium glutamate, 10% glycerol, 1 mM BME). Peak fractions were assessed by SDS–PAGE and subsequently pooled, concentrated, aliquoted, and flash-frozen in liquid nitrogen and stored at −80°C.

### Phase separation studies of human Orc1^IDR^

The His6-MBP tag was removed from purified Orc1^IDR^ and Orc1^IDR-PM^ by TEV cleavage (at a 1/10 mass ratio) for 1 hr at RT before preparing samples containing stoichiometric amounts (final concentrations = 0.5 µM, 1 µM, 2 µM, and 5 µM) of 60 basepair Cy5-labeled duplex DNA in Assay Buffer (50mM HEPES, 150mM K-Glut, 10% Glycerol). The protein-DNA mixture was then added to an imaging plate (Ibidi, µ-Plate, Cat. #89626), incubated on ice for 15 min, and then centrifuged for 3 min at 20 g to remove bubbles. The samples were imaged at RT using a Nikon Ti2E microscope equipped with a Yokogawa CSU X1 spinning disk with a 60× oil immersion objective, an appropriate filter set, and 641 nm laser excitation (2% laser power). Z-stacks were collected at 0.5 µm steps through the protein networks. Images were analyzed in FIJI by first generating maximum intensity projections and then subtracting background signal calculated from a negative control (buffer only).

### Fluorescence polarization DNA binding assays

Serial dilutions of Orc1^IDR^ were made in Sizing Buffer (50 mM HEPES pH 7.5, 10% glycerol, 300 mM K-Glut, and 1 mM BME), and an equal volume of 20 nM FITC-dsDNA prepared in Dilution Buffer (50 mM HEPES pH 7.5, 10% glycerol, and 1 mM BME) was added to each. Reactions were incubated for 30 min at room temperature and then transferred to a 384-well black-bottom multi-well plate (Corning, Cat. #3820). Fluorescence polarization measurements were made using a CLARIOstar BMG LABTECH plate reader. All intensities were background-corrected first for a buffer blank, and then polarizations were calculated for a fluorophore-only sample. The data were plotted and fit with a nonlinear regression curve for specific binding with Hill slope using GraphPad Prism to calculate the apparent dissociation constant (K_d_). The K_d_ reported represents the average and standard deviation of three independent experiments.

### Analysis of genomic binding profiles by CUT&RUN

Cells were transfected as described above. Twenty-four hours after transfection, cells were detached using trypsin (Gibco, Cat. #25300-062), and enzymatic activity was quenched by the addition of complete DMEM. Cells were collected by centrifugation at 600 × g for 3 minutes, resuspended in phosphate-buffered saline (PBS; Gibco, Cat. #10010-23) and washed once by an additional centrifugation step.CUT&RUN was performed using the CUT&RUN Assay Kit (Active Motif, Cat. #53180) according to the manufacturer’s instructions. Briefly, concanavalin A-coated magnetic beads were prepared by washing twice with the 1x binding buffer and then kept on ice until use. Cells were washed with wash buffer and incubated in nuclei isolation buffer to release nuclei. Isolated nuclei were washed and incubated with the prepared concanavalin A beads to immobilize the nuclei. Samples were then incubated overnight at 4°C with rotation in the presence of 1 μg of either anti-FLAG antibody (EpiCypher, CUT&RUN grade, Cat. #13-203) for all Orc1 and Cdt1 constructs or anti-H3K4me3 antibody provided by the manufacturer for H3K4me3 experiments. The following day, samples were washed and incubated with permeabilization buffer containing MNase. Calcium chloride (CaCl₂) was added to activate MNase digestion, and samples were incubated for 2 hours at 4°C. Digestion was terminated according to the manufacturer’s protocol; chromatin fragments were released by heat treatment, and the DNA was purified and eluted in 55 µl of water.

Samples were submitted to the UT Southwestern Genomics Core for library preparation and next-generation sequencing. Sample concentration was measured using the PicoGreen dsDNA Quantitation Assay Kit (Invitrogen, Cat. #P7589) combined with a fluorescence-based microplate reader (PerkinElmer Victor X3 Multilabel Reader, Model 2030). The KAPA HyperPlus Library Preparation Kit (Roche/KAPA, Cat. #07962428001) was used to prepare DNA libraries. The workflow included DNA fragment end repair, A-tailing, adapter and index barcode ligation, and two rounds of AMPure XP bead size selection. The adapter-ligated libraries were then PCR-amplified to generate the final sequencing libraries, followed by two rounds of AMPure XP bead purification before the final library quality and quantity assessments. Library quantity was measured using the PicoGreen method, and library quality was evaluated using the Agilent 2100 Bioanalyzer with the DNA High Sensitivity Kit (Agilent Technologies, Catalog #5067-4626). All libraries were required to meet quality control standards before sequencing could proceed. The libraries were sequenced on either an Illumina NextSeq platform or an Illumina NovaSeq platform using a paired-end sequencing protocol, following the manufacturer’s instructions.

### CUT&RUN data analysis pipeline

All genomics data, including ATAC-seq data obtained from GSM5128944, were processed using the Galaxy platform. Raw sequencing reads were first assessed for quality using FastQC and subsequently trimmed with Trim Galore! using Illumina Universal Adapters. Reads with a Phred quality score below 30 were removed and reads shorter than 15 bp after trimming were discarded. The resulting high-quality reads were aligned to the hg38 canonical female reference genome using Bowtie2 in paired-end, very-sensitive, end-to-end mode, with a maximum fragment length of 1000 bp. Following alignment, BAM files were filtered to retain properly paired reads with a mapping quality score of at least 30, while reads aligning to the mitochondrial chromosome (chrM) were excluded. PCR duplicates were subsequently identified and removed using MarkDuplicates. Peaks were then identified using MACS2 callpeak with a q-value threshold of < 0.2. For downstream visualization and quantitative analysis, genome-wide coverage files were generated in BigWig format using BamCoverage with a 50-bp bin size and visualized using IGV (version 2.18.2). Signal enrichment across genomic regions of interest was evaluated by generating heatmaps and average signal profiles using computeMatrix and plotHeatmap. Finally, genome-wide signal correlations between datasets were assessed using BigWigSummary with a bin size of 5,000 bp, followed by visualization with plotCorrelation.

### Computational analysis of protein disordered regions

Metapredict was used to predict disordered regions (*78*). A custom Python script was used to calculate sequence complexity and charged residue parameters, including the position of positively (K,R) and negatively charged amino acids (D,E) and their distribution across the sequence (Kappa, (*79*)).

## ACKNOWLEDGMENTS

We thank past and present members of the Parker Lab for helpful discussion and technical advice. This work was supported by the National Science Foundation (NSF 2308642, to M.W.P.), the Cancer Prevention and Research Institute of Texas (CPRIT RR200070, to M.W.P.), and the Welch Foundation (V-I-0004-20230731, to M.W.P.). M.W.P is the Cecil H. and Ida Green Endowed Scholar in Biomedical Computational Science.

## DATA AVAILABILITY

All raw and analyzed data presented in this paper are publicly available through our lab’s Data Dryad repository.

---

**Figure S1:**
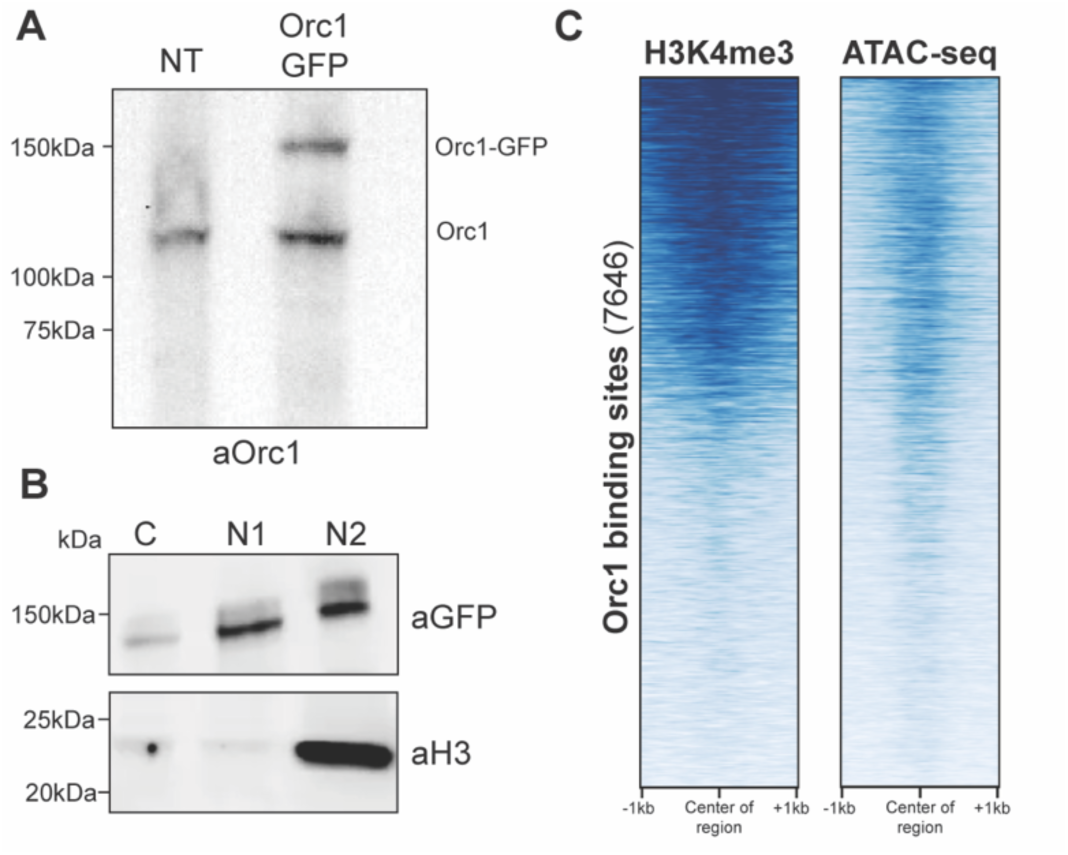
Analysis of Orc1 chromatin association. (A) Analysis of protein expression levels by Western blot of HeLa cell lysate. The endogenous Orc1 and transgene-expressed Orc1 are indicated with “Orc1” and “Orc1-GFP”, respectively. (B) Cellular fractionation studies to determine the cellular localization and chromatin association of Orc1. Histone H3 was analyzed as a chromatin-associated control. Fractions include: “C” = cytoplasm, “N1” = nuclear soluble, and “N2” = nuclear bound. (C) Heatmap representation of sequencing read intensity of H3K4me3 (first panel, data generated in this study) and open regions (ATAC-seq, GEO = GSM5128944) across Orc1 peaks (+/− 1 kb).

**Figure S2:**
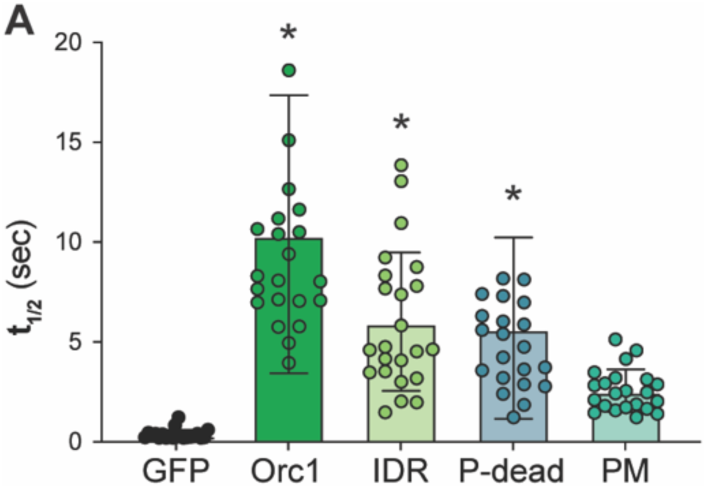
FRAP analysis of chromatin binding in HEK293 cells. (A) Half-time of recovery calculated from FRAP analysis of GFP, Orc1, Orc1^IDR^, Orc1^IDR-P-dead^, and Orc1^IDR-PM^ in HEK293 cells. Significance calculated using one-way ANOVA test compared to GFP (* = p < 0.05).

**Figure S3:**
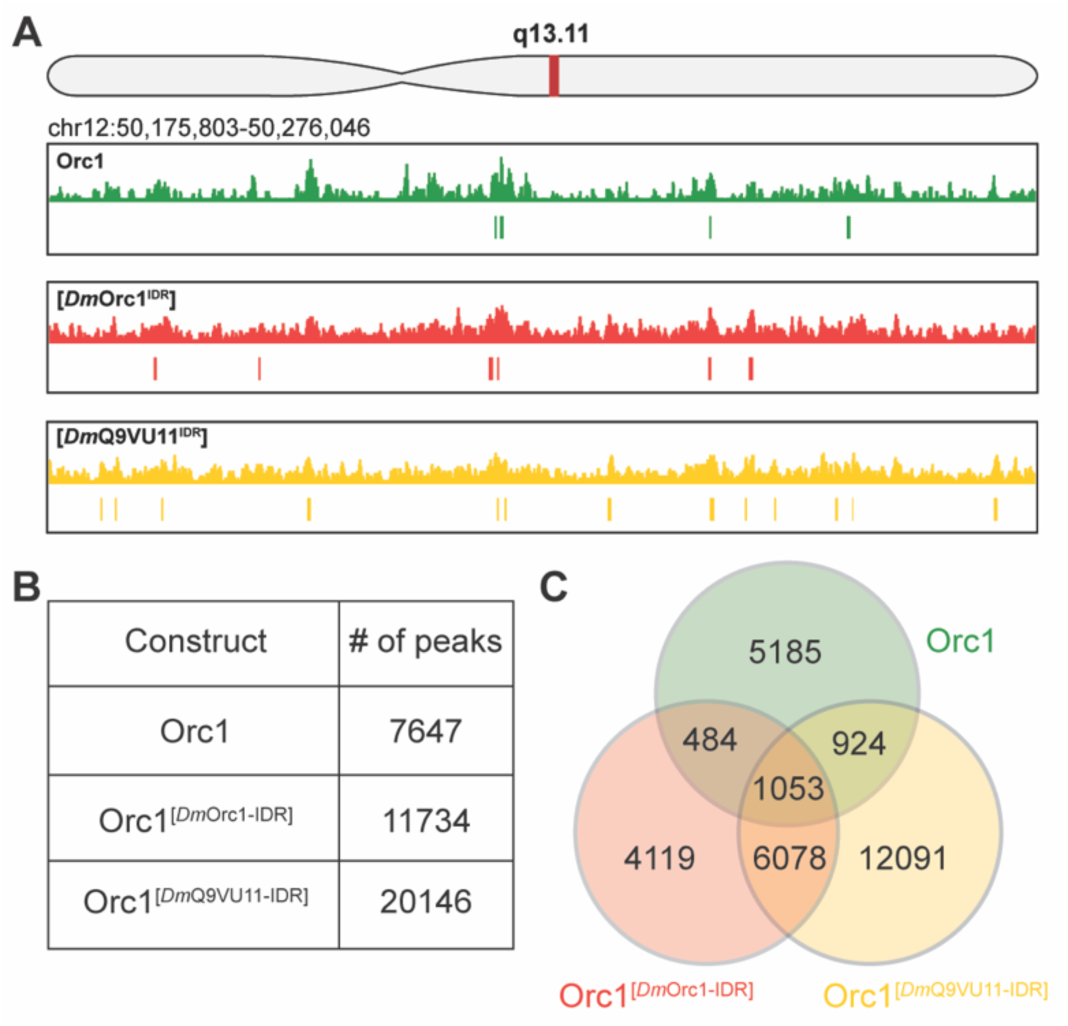
Comparison of Orc1 chimeras’ genome binding profiles. (A) Genome browser view of a 100 kb region showing the CUT&RUN sequencing read intensity from Orc1 (top panel) and two Orc1 chimeras (middle and bottom panel). Peaks are indicated with a hash mark. (B) Number of MACS2-called peaks for Orc1 and Orc1 chimeras. (C) Venn diagram representing overlapping peaks between Orc1 (green circle) and Orc1 chimeras (red = Orc1^[*Dm*Orc1-IDR]^ and yellow = Orc1^[*Dm*Q9VU11-IDR]^).

**Figure S4:**
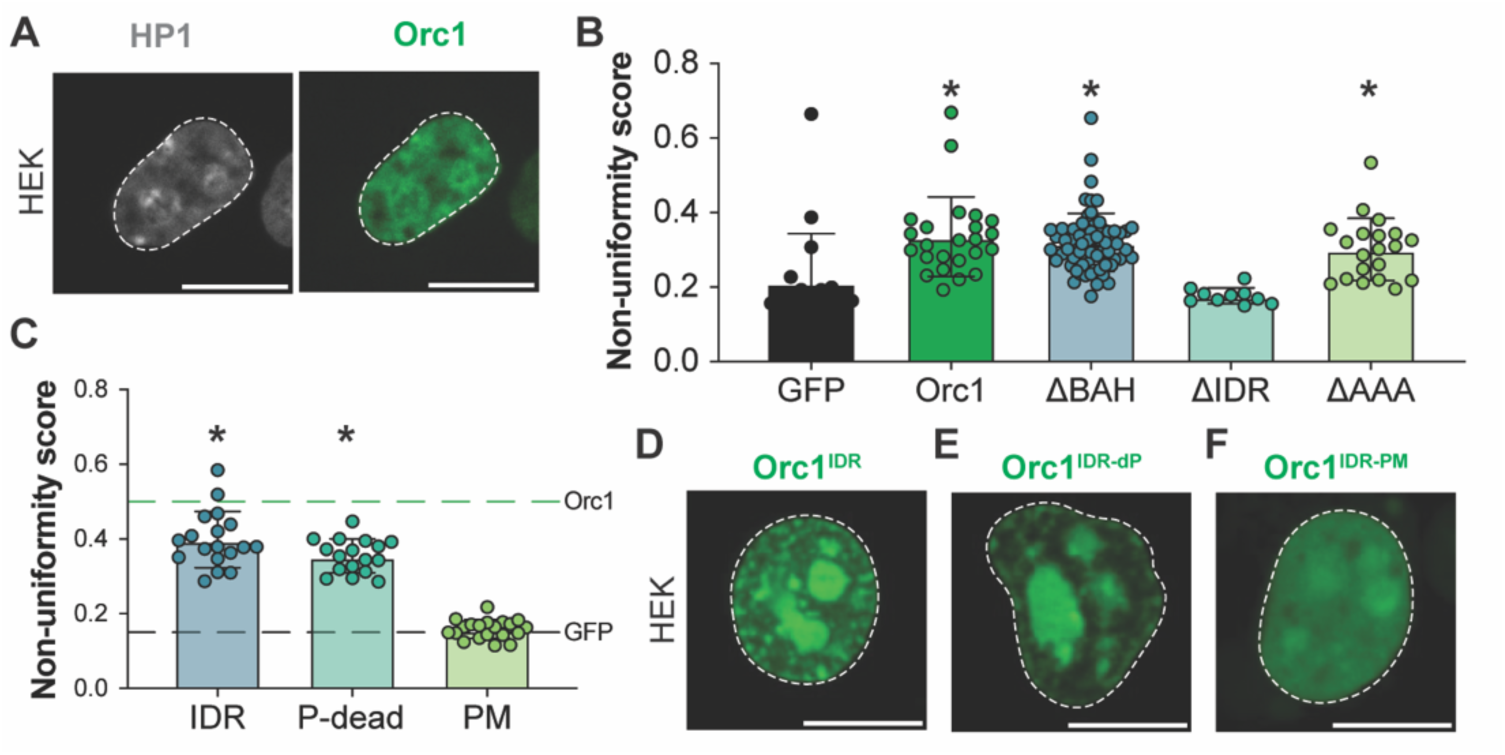
Role of the Orc1 IDR in heterochromatin partitioning. (A) Confocal fluorescence image showing the nuclear distribution of mCherry-tagged Heterochromatin protein 1*a* (Hp1*a*) and GFP-tagged human Orc1 in HEK293 cells. Scale bar = 10 µm. (B) Nuclear non-uniformity (n.u.) of GFP, Orc1, Orc1^ΔBAH^, Orc1^ΔIDR^, and Orc1^ΔAAA^ in HeLa cells. Significance was established by a one-way ANOVA test with comparison to GFP (* = p < 0.05). (C) Nuclear non-uniformity (n.u.) of Orc1^IDR^, Orc1^IDR-P-dead^, and Orc1^IDR-PM^ in HEK293 cells. Significance was established by a one-way ANOVA test with comparison to GFP (* = p < 0.05). (D-F) Confocal fluorescence images showing the nuclear distribution of GFP-tagged (D) Orc1^IDR^, (E) Orc1^IDR-P-dead^, and (F) Orc1^IDR-PM^ in HEK293 cells. Scale bar = 10 µm.

**Figure S5:**
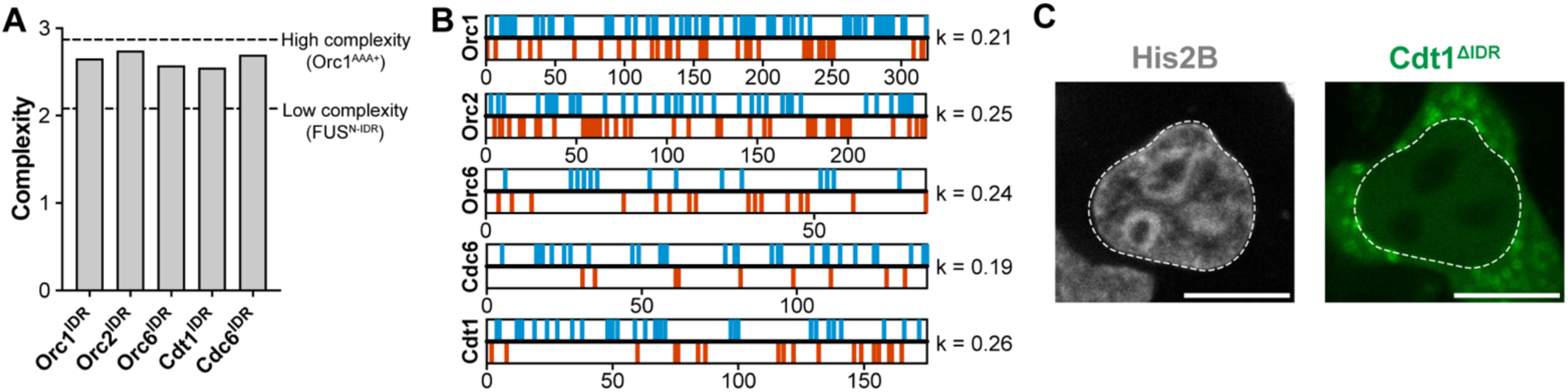
Sequence features of licensing factor IDRs. (A) Sequence complexity of licensing factor IDRs. The Orc1 AAA+ domain (amino acids 526-734) is used as a representative “high-complexity” sequence, and the Fus N-terminal IDR (amino acids 1-287) is used as a representative “low-complexity” sequence. (B) Distribution of positively charged residues (Arg and Lys, blue) and negatively charged residues (Asp and Glu, red) in licensing factor IDRs. Charged residue distribution was quantified using kappa (k, (*79*)) and the score for each is indicated. (C) Representative image of Cdt1^ΔIDR^ in HeLa cells. Scale bar = 10 μm.

## BIBLIOGRAPHY

1. T. M. Weaver, E. A. Morrison, C. A. Musselman, Reading More than Histones: The Prevalence of Nucleic Acid Binding among Reader Domains. Molecules 23, 2614 (2018).

2. M. Lachner, D. O’Carroll, S. Rea, K. Mechtler, T. Jenuwein, Methylation of histone H3 lysine 9 creates a binding site for HP1 proteins. Nature 410, 116–120 (2001).

3. M. Ninova, K. Fejes Tóth, A. A. Aravin, The control of gene expression and cell identity by H3K9 trimethylation. Development 146, dev181180 (2019).

4. C. O. Pabo, R. T. Sauer, TRANSCRIPTION FACTORS: Structural Families and Principles of DNA Recognition. Annu. Rev. Biochem. 61, 1053–1095 (1992).

5. R. Li, L. Yang, E. Fouts, M. R. Botchan, Site-specific DNA-binding Proteins Important for Replication and Transcription Have Multiple Activities. Cold Spring Harbor Symposia on Quantitative Biology 58, 403–413 (1993).

6. M. I. Aladjem, C. E. Redon, Order from clutter: selective interactions at mammalian replication origins. Nat Rev Genet 18, 101–116 (2017).

7. P. J. Mitchell, R. Tjian, Transcriptional Regulation in Mammalian Cells by Sequence-Specific DNA Binding Proteins. Science 245, 371–378 (1989).

8. R. Rohs, X. Jin, S. M. West, R. Joshi, B. Honig, R. S. Mann, Origins of Specificity in Protein-DNA Recognition. Annu. Rev. Biochem. 79, 233–269 (2010).

9. C. Wolberger, How structural biology transformed studies of transcription regulation. Journal of Biological Chemistry 296, 100741 (2021).

10. M. Slattery, T. Zhou, L. Yang, A. C. Dantas Machado, R. Gordân, R. Rohs, Absence of a simple code: how transcription factors read the genome. Trends in Biochemical Sciences 39, 381–399 (2014).

11. J. F. Kribelbauer, C. Rastogi, H. J. Bussemaker, R. S. Mann, Low-Affinity Binding Sites and the Transcription Factor Specificity Paradox in Eukaryotes. Annu. Rev. Cell Dev. Biol. 35, 357–379 (2019).

12. J. Burdach, A. P. W. Funnell, K. S. Mak, C. M. Artuz, B. Wienert, W. F. Lim, L. Y. Tan, R. C. M. Pearson, M. Crossley, Regions outside the DNA-binding domain are critical for proper in vivo specificity of an archetypal zinc finger transcription factor. Nucleic Acids Research 42, 276–289 (2014).

13. S. Inukai, K. H. Kock, M. L. Bulyk, Transcription factor–DNA binding: beyond binding site motifs. Current Opinion in Genetics & Development 43, 110–119 (2017).

14. J. Liu, D. K. Kumar, B. Hurieva, F. Jonas, N. Barkai, Engineering intrinsically disordered regions for guiding genome navigation. Molecular Cell 86, 851–867.e5 (2026).

15. S. Brodsky, T. Jana, K. Mittelman, M. Chapal, D. K. Kumar, M. Carmi, N. Barkai, Intrinsically Disordered Regions Direct Transcription Factor In Vivo Binding Specificity. Molecular Cell 79, 459–471.e4 (2020).

16. A. A. Abidi, C. Cattoglio, N. N. Tang, V. B. Fan, G. M. Dailey, A. D. Hay, P. Kunamaneni, D. E. Milkie, X. Darzacq, E. Betzig, R. Tjian, T. G. W. Graham, Unstructured transcription factor interactions enable emergent specificity. Science 392, eaeb6487 (2026).

17. S. Brodsky, T. Jana, N. Barkai, Order through disorder: The role of intrinsically disordered regions in transcription factor binding specificity. Current Opinion in Structural Biology 71, 110–115 (2021).

18. F. Bleichert, M. R. Botchan, J. M. Berger, Mechanisms for initiating cellular DNA replication. Science 355, eaah6317 (2017).

19. S. P. Bell, A. Dutta, DNA Replication in Eukaryotic Cells. Annu. Rev. Biochem. 71, 333–374 (2002).

20. M. Fragkos, O. Ganier, P. Coulombe, M. Méchali, DNA replication origin activation in space and time. Nat Rev Mol Cell Biol 16, 360–374 (2015).

21. H. Li, M. E. O’Donnell, The Eukaryotic CMG Helicase at the Replication Fork: Emerging Architecture Reveals an Unexpected Mechanism. BioEssays 40, 1700208 (2018).

22. S. P. Bell, B. Stillman, ATP-dependent recognition of eukaryotic origins of DNA replication by a multiprotein complex. Nature 357, 128–134 (1992).

23. J.-K. Lee, K.-Y. Moon, Y. Jiang, J. Hurwitz, The *Schizosaccharomyces pombe* origin recognition complex interacts with multiple AT-rich regions of the replication origin DNA by means of the AT-hook domains of the spOrc4 protein. Proc. Natl. Acad. Sci. U.S.A. 98, 13589–13594 (2001).

24. R.-Y. Chuang, T. J. Kelly, The fission yeast homologue of Orc4p binds to replication origin DNA via multiple AT-hooks. Proc. Natl. Acad. Sci. U.S.A. 96, 2656–2661 (1999).

25. S. Vashee, C. Cvetic, W. Lu, P. Simancek, T. J. Kelly, J. C. Walter, Sequence-independent DNA binding and replication initiation by the human origin recognition complex. Genes Dev. 17, 1894–1908 (2003).

26. H. K. MacAlpine, R. Gordân, S. K. Powell, A. J. Hartemink, D. M. MacAlpine, *Drosophila* ORC localizes to open chromatin and marks sites of cohesin complex loading. Genome Res. 20, 201–211 (2010).

27. G. I. Dellino, D. Cittaro, R. Piccioni, L. Luzi, S. Banfi, S. Segalla, M. Cesaroni, R. Mendoza-Maldonado, M. Giacca, P. G. Pelicci, Genome-wide mapping of human DNA-replication origins: Levels of transcription at ORC1 sites regulate origin selection and replication timing. Genome Res. 23, 1–11 (2013).

28. K. Siddiqui, B. Stillman, ATP-dependent Assembly of the Human Origin Recognition Complex. Journal of Biological Chemistry 282, 32370–32383 (2007).

29. S. Ohta, Y. Tatsumi, M. Fujita, T. Tsurimoto, C. Obuse, The ORC1 Cycle in Human Cells. Journal of Biological Chemistry 278, 41535–41540 (2003).

30. N. Xu, Y. You, C. Liu, M. Balasov, L. T. Lun, Y. Geng, C. P. Fung, H. Miao, H. Tian, T. T. Choy, X. Shi, Z. Fan, B. Zhou, K. Akhmetova, R. U. Din, H. Yang, Q. Hao, P. Qian, I. Chesnokov, G. Zhu, Structural basis of DNA replication origin recognition by human Orc6 protein binding with DNA. Nucleic Acids Research 48, 11146–11161 (2020).

31. A. J. Kuo, J. Song, P. Cheung, S. Ishibe-Murakami, S. Yamazoe, J. K. Chen, D. J. Patel, O. Gozani, The BAH domain of ORC1 links H4K20me2 to DNA replication licensing and Meier– Gorlin syndrome. Nature 484, 115–119 (2012).

32. F. Bleichert, M. R. Botchan, J. M. Berger, Crystal structure of the eukaryotic origin recognition complex. Nature 519, 321–326 (2015).

33. O. A. Adiji, B. S. McConnell, M. W. Parker, The origin recognition complex requires chromatin tethering by a hypervariable intrinsically disordered region that is functionally conserved from sponge to man. Nucleic Acids Research 52, 4344–4360 (2024).

34. G. Xouri, A. Squire, M. Dimaki, B. Geverts, P. J. Verveer, S. Taraviras, H. Nishitani, A. B. Houtsmuller, P. I. H. Bastiaens, Z. Lygerou, Cdt1 associates dynamically with chromatin throughout G1 and recruits Geminin onto chromatin. EMBO J 26, 1303–1314 (2007).

35. S. G. Prasanth, Z. Shen, K. V. Prasanth, B. Stillman, Human origin recognition complex is essential for HP1 binding to chromatin and heterochromatin organization. Proc. Natl. Acad. Sci. U.S.A. 107, 15093–15098 (2010).

36. F. Mueller, T. S. Karpova, D. Mazza, J. G. McNally, “Monitoring Dynamic Binding of Chromatin Proteins In Vivo by Fluorescence Recovery After Photobleaching” in Chromatin Remodeling, R. H. Morse, Ed. (Humana Press, Totowa, NJ, 2012; https://link.springer.com/10.1007/978-1-61779-477-3_11)vol. 833 of *Methods in Molecular Biology*, pp. 153–176.

37. N. Kara, M. Hossain, S. G. Prasanth, B. Stillman, Orc1 Binding to Mitotic Chromosomes Precedes Spatial Patterning during G1 Phase and Assembly of the Origin Recognition Complex in Human Cells. Journal of Biological Chemistry 290, 12355–12369 (2015).

38. L. S. Bicknell, E. M. H. F. Bongers, A. Leitch, S. Brown, J. Schoots, M. E. Harley, S. Aftimos, J. Y. Al-Aama, M. Bober, P. A. J. Brown, H. Van Bokhoven, J. Dean, A. Y. Edrees, M. Feingold, A. Fryer, L. H. Hoefsloot, N. Kau, N. V. A. M. Knoers, J. MacKenzie, J. M. Opitz, P. Sarda, A. Ross, I. K. Temple, A. Toutain, C. A. Wise, M. Wright, A. P. Jackson, Mutations in the pre-replication complex cause Meier-Gorlin syndrome. Nat Genet 43, 356–359 (2011).

39. P. J. Skene, S. Henikoff, An efficient targeted nuclease strategy for high-resolution mapping of DNA binding sites. eLife 6, e21856 (2017).

40. O. A. Adiji, I. Leonovich, M. W. Parker, Phosphorylation alters the bulk chemical properties of Orc1 to tune DNA binding, phase separation, and heterochromatin partitioning. Biochemistry [Preprint] (2026). 10.64898/2026.08.17.745304.

41. D. T. S. Pak, M. Pflumm, I. Chesnokov, D. W. Huang, R. Kellum, J. Marr, P. Romanowski, M. R. Botchan, Association of the Origin Recognition Complex with Heterochromatin and HP1 in Higher Eukaryotes. Cell 91, 311–323 (1997).

42. Y. Luo, S. Rajshekar, H. Kumar, J. M. Berger, G. H. Karpen, M. R. Botchan, ORC binding to Heterochromatin Protein 1 through intrinsically disordered regions is required for heterochromatin structure and function. Biochemistry [Preprint] (2026). 10.64898/2026.07.17.738698.

43. A. G. Larson, D. Elnatan, M. M. Keenen, M. J. Trnka, J. B. Johnston, A. L. Burlingame, D. A. Agard, S. Redding, G. J. Narlikar, Liquid droplet formation by HP1α suggests a role for phase separation in heterochromatin. Nature 547, 236–240 (2017).

44. A. R. Strom, A. V. Emelyanov, M. Mir, D. V. Fyodorov, X. Darzacq, G. H. Karpen, Phase separation drives heterochromatin domain formation. Nature 547, 241–245 (2017).

45. M. W. Parker, M. Bell, M. Mir, J. A. Kao, X. Darzacq, M. R. Botchan, J. M. Berger, A new class of disordered elements controls DNA replication through initiator self-assembly. eLife 8, e48562 (2019).

46. M. Hossain, K. Bhalla, B. Stillman, Multiple, short protein binding motifs in ORC1 and CDC6 control the initiation of DNA replication. Molecular Cell 81, 1951–1969.e6 (2021).

47. H. Nishitani, S. Taraviras, Z. Lygerou, T. Nishimoto, The Human Licensing Factor for DNA Replication Cdt1 Accumulates in G1 and Is Destabilized after Initiation of S-phase. Journal of Biological Chemistry 276, 44905–44911 (2001).

48. D. Coverley, C. Pelizon, S. Trewick, R. A. Laskey, Chromatin-bound Cdc6 persists in S and G2 phases in human cells, while soluble Cdc6 is destroyed in a cyclin A-cdk2 dependent process. Journal of Cell Science 113, 1929–1938 (2000).

49. W. Jiang, N. J. Wells, T. Hunter, Multistep regulation of DNA replication by Cdk phosphorylation of HsCdc6. Proc. Natl. Acad. Sci. U.S.A. 96, 6193–6198 (1999).

50. K. Yanagi, T. Mizuno, Z. You, F. Hanaoka, Mouse Geminin Inhibits Not Only Cdt1-MCM6 Interactions but Also a Novel Intrinsic Cdt1 DNA Binding Activity. Journal of Biological Chemistry 277, 40871–40880 (2002).

51. N. Li, W. H. Lam, Y. Zhai, J. Cheng, E. Cheng, Y. Zhao, N. Gao, B.-K. Tye, Structure of the origin recognition complex bound to DNA replication origin. Nature 559, 217–222 (2018).

52. D. Kong, M. L. DePamphilis, Site-Specific DNA Binding of the *Schizosaccharomyces pombe* Origin Recognition Complex Is Determined by the Orc4 Subunit. Molecular and Cellular Biology 21, 8095–8103 (2001).

53. B. Miotto, Z. Ji, K. Struhl, Selectivity of ORC binding sites and the relation to replication timing, fragile sites, and deletions in cancers. Proc. Natl. Acad. Sci. U.S.A. 113 (2016).

54. D. Remus, E. L. Beall, M. R. Botchan, DNA topology, not DNA sequence, is a critical determinant for Drosophila ORC–DNA binding. EMBO J 23, 897–907 (2004).

55. K. Noguchi, A. Vassilev, S. Ghosh, J. L. Yates, M. L. DePamphilis, The BAH domain facilitates the ability of human Orc1 protein to activate replication origins in vivo. EMBO J 25, 5372–5382 (2006).

56. M. W. Parker, J. A. Kao, A. Huang, J. M. Berger, M. R. Botchan, Molecular determinants of phase separation for Drosophila DNA replication licensing factors. eLife 10, e70535 (2021).

57. O. Hunker, F. Bleichert, AlphaFold-guided phylogenetic analyses suggest surprising heterogeneity in metazoan replication origin licensing mechanisms. EMBO J 45, 13 (2026).

58. S. Jorgensen, G. Schotta, C. S. Sorensen, Histone H4 Lysine 20 methylation: key player in epigenetic regulation of genomic integrity. Nucleic Acids Research 41, 2797–2806 (2013).

59. F. Bleichert, A. Leitner, R. Aebersold, M. R. Botchan, J. M. Berger, Conformational control and DNA-binding mechanism of the metazoan origin recognition complex. Proc. Natl. Acad. Sci. U.S.A. 115 (2018).

60. J. A. Belsky, H. K. MacAlpine, Y. Lubelsky, A. J. Hartemink, D. M. MacAlpine, Genome-wide chromatin footprinting reveals changes in replication origin architecture induced by pre-RC assembly. Genes Dev. 29, 212–224 (2015).

61. D. E. Schones, K. Cui, S. Cuddapah, T.-Y. Roh, A. Barski, Z. Wang, G. Wei, K. Zhao, Dynamic Regulation of Nucleosome Positioning in the Human Genome. Cell 132, 887–898 (2008).

62. S. Henikoff, Nucleosome destabilization in the epigenetic regulation of gene expression. Nat Rev Genet 9, 15–26 (2008).

63. J. C. Bell, D. Jukam, N. A. Teran, V. I. Risca, O. K. Smith, W. L. Johnson, J. M. Skotheim, W. J. Greenleaf, A. F. Straight, Chromatin-associated RNA sequencing (ChAR-seq) maps genome-wide RNA-to-DNA contacts. eLife 7, e27024 (2018).

64. J. Norseen, A. Thomae, V. Sridharan, A. Aiyar, A. Schepers, P. M. Lieberman, RNA-dependent recruitment of the origin recognition complex. EMBO J 27, 3024–3035 (2008).

65. A. Eladl, Y. Yamaoki, S. Hoshina, H. Horinouchi, K. Kondo, S. Waga, T. Nagata, M. Katahira, Investigation of the Interaction of Human Origin Recognition Complex Subunit 1 with G-Quadruplex DNAs of Human c-myc Promoter and Telomere Regions. IJMS 22, 3481 (2021).

66. S. Ummarino, L. Poluben, A. K. Ebralidze, I. Autiero, L. Rinaldi, T. Paniza, M. Deshpande, N. H. Mandel, J. D. Lee, Y. Zhang, M. A. Bassal, B. Budnik, B. Q. Trinh, S. P. Balk, R. Flaumenhaft, J. Gerhardt, S. M. Mirkin, D. G. Tenen, A. Di Ruscio, RNAs anchoring replication complex control initiation and firing of DNA replication. Nat Commun, doi: 10.1038/s41467-026-73478-2 (2026).

67. C. A. Seller, P. H. O’Farrell, Rif1 prolongs the embryonic S phase at the Drosophila mid-blastula transition. PLoS Biol 16, e2005687 (2018).

68. R. Sukackaite, D. Cornacchia, M. R. Jensen, P. J. Mas, M. Blackledge, E. Enervald, G. Duan, T. Auchynnikava, M. Köhn, D. J. Hart, S. B. C. Buonomo, Mouse Rif1 is a regulatory subunit of protein phosphatase 1 (PP1). Sci Rep 7, 2119 (2017).

69. S. Hiraga, G. M. Alvino, F. Chang, H. Lian, A. Sridhar, T. Kubota, B. J. Brewer, M. Weinreich, M. K. Raghuraman, A. D. Donaldson, Rif1 controls DNA replication by directing Protein Phosphatase 1 to reverse Cdc7-mediated phosphorylation of the MCM complex. Genes Dev. 28, 372–383 (2014).

70. M. Bollen, W. Peti, M. J. Ragusa, M. Beullens, The extended PP1 toolkit: designed to create specificity. Trends in Biochemical Sciences 35, 450–458 (2010).

71. P. Vagnarelli, S. Ribeiro, L. Sennels, L. Sanchez-Pulido, F. de Lima Alves, T. Verheyen, D. A. Kelly, C. P. Ponting, J. Rappsilber, W. C. Earnshaw, Repo-Man Coordinates Chromosomal Reorganization with Nuclear Envelope Reassembly during Mitotic Exit. Developmental Cell 21, 328–342 (2011).

72. I. J. De Castro, J. Budzak, M. L. Di Giacinto, L. Ligammari, E. Gokhan, C. Spanos, D. Moralli, C. Richardson, J. I. De Las Heras, S. Salatino, E. C. Schirmer, K. S. Ullman, W. A. Bickmore, C. Green, J. Rappsilber, S. Lamble, M. W. Goldberg, V. Vinciotti, P. Vagnarelli, Repo-Man/PP1 regulates heterochromatin formation in interphase. Nat Commun 8, 14048 (2017).

73. H. B. Landsverk, F. Mora-Bermúdez, O. J. B. Landsverk, G. Hasvold, S. Naderi, O. Bakke, J. Ellenberg, P. Collas, R. G. Syljuåsen, T. Küntziger, The protein phosphatase 1 regulator PNUTS is a new component of the DNA damage response. EMBO Rep 11, 868–875 (2010).

74. D. K. Kumar, F. Jonas, T. Jana, S. Brodsky, M. Carmi, N. Barkai, Complementary strategies for directing in vivo transcription factor binding through DNA binding domains and intrinsically disordered regions. Molecular Cell 83, 1462–1473.e5 (2023).

75. Z. Peng, J. Yan, X. Fan, M. J. Mizianty, B. Xue, K. Wang, G. Hu, V. N. Uversky, L. Kurgan, Exceptionally abundant exceptions: comprehensive characterization of intrinsic disorder in all domains of life. Cell. Mol. Life Sci. 72, 137–151 (2015).

76. J. J. Ward, J. S. Sodhi, L. J. McGuffin, B. F. Buxton, D. T. Jones, Prediction and Functional Analysis of Native Disorder in Proteins from the Three Kingdoms of Life. Journal of Molecular Biology 337, 635–645 (2004).

77. F. L. Goerner, T. Duong, R. J. Stafford, G. D. Clarke, A comparison of five standard methods for evaluating image intensity uniformity in partially parallel imaging MRI: Measuring uniformity in MRI with partially parallel imaging. Med. Phys. 40, 082302 (2013).

78. J. M. Lotthammer, J. Hernández-García, D. Griffith, D. Weijers, A. S. Holehouse, R. J. Emenecker, Metapredict enables accurate disorder prediction across the Tree of Life. Bioinformatics [Preprint] (2024). 10.1101/2024.11.05.622168.

79. R. K. Das, R. V. Pappu, Conformations of intrinsically disordered proteins are influenced by linear sequence distributions of oppositely charged residues. Proc. Natl. Acad. Sci. U.S.A. 110, 13392–13397 (2013).

